# A stratified benchmark reveals distinct short- and long-read performance gradients within difficult genomic regions

**DOI:** 10.64898/2026.11.26.769538

**Authors:** Xiwen Zhang, Yichen Henry Liu, Yujie Wang, Xin Maizie Zhou

**Affiliations:** Department of Biological Sciences, Vanderbilt University, 37235 Nashville, USA; Department of Computer Science, Vanderbilt University, 37235 Nashville, USA; Department of Biomedical Informatics, Harvard Medical School, 02115 Boston, USA; Department of Biomedical Engineering, Vanderbilt University, 37235 Nashville, USA

**Author notes:** Equal contributor (Xiwen Zhang, Yichen Henry Liu, Yujie Wang). Full list of author information is available at the end of the article.

**Keywords:** Variant calling, Whole genome sequencing, Genome stratification, Low mappability regions, Segmental duplication

## Abstract

Accurate detection of small variants underpins population, functional and clinical genomics, yet short-read sequencing remains unreliable in repetitive and duplicated regions of the human genome, where long reads provide substantially greater sequence context. Although this disparity is well documented, aggregate accuracy metrics delineate where short- and long-read callsets diverge but not the mechanisms that give rise to these differences. Here, we present a region-stratified benchmark of single-nucleotide polymorphism (SNP) and insertion–deletion (INDEL) detection from Illumina short-read and PacBio HiFi long-read sequencing of the Genome in a Bottle reference sample HG002, evaluating multiple variant callers across genomic contexts, sequencing depths and INDEL lengths. Both platforms achieved near-complete accuracy in high-confidence regions, whereas short-read performance deteriorated in low-mappability regions and segmental duplications, a deficit that persisted with increasing sequencing depth. Within these regions, short-read recall declined toward the interiors of long tracts, with increasing sequence identity to the closest paralog and with greater distance from paralogous sequence variants, and these declines coincided with a pronounced excess of ambiguously placed reads. Together, these findings implicate ambiguous read placement as the predominant source of the short-read deficit, demonstrate that its magnitude can be anticipated from the length and paralog similarity of the surrounding sequence, and thereby delineate the genomic contexts in which long-read sequencing is most warranted.

## Introduction

Small genetic variants, including single-nucleotide variants (SNVs) and short insertions and deletions (in-dels), constitute the most abundant form of sequence variation in the human genome [1, 2]. Although most small variants are functionally neutral, a subset alters protein-coding sequences, gene regulation, RNA processing, or chromatin organization and contributes to Mendelian disorders, complex traits, and inter-individual phenotypic diversity [3]. Accurate small-variant detection is therefore foundational to population genetics, functional genomics, clinical sequencing, and studies of human disease.

Short-read sequencing has established the technical and computational foundation of contemporary small-variant analysis [4]. The high per-base accuracy, throughput, and cost efficiency of platforms such as Illumina, together with mature alignment, variant-calling, and quality-calibration frameworks, enable highly accurate SNV and short-indel detection across much of the genome [5–7]. In regions that are uniquely mappable and adequately represented by a linear reference, state-of-the-art short-read pipelines can achieve near-saturated performance against established bench-mark callsets [8]. Nevertheless, this performance is un-even across genomic contexts. Short reads frequently lack sufficient sequence context to resolve paralogous sequences, segmental duplications, tandem repeats, homopolymers, highly polymorphic loci, and regions containing nearby or nested variants [9, 10]. In these settings, ambiguous mapping, reference bias, incomplete local haplotype reconstruction, and inconsistent representation of complex alleles can reduce sensitivity and increase false-positive calls. These limitations are particularly consequential because difficult-to-analyse regions are enriched for duplicated genes, immune-related loci, medically relevant repeat expansions, and other functionally important sequences [11, 12].

Long-read sequencing offers a complementary view of small genetic variation [13, 14]. Pacific Biosciences circular consensus sequencing produces highly accurate long reads [15, 16], whereas Oxford Nanopore Technologies can generate reads spanning tens to hundreds of kilobases and directly capture additional molecular features [17, 18]. Their extended sequence context improves read placement, haplotype resolution, and the interpretation of variants embedded within repetitive or structurally complex regions [16]. Improvements in sequencing chemistry, consensus generation, basecalling, alignment, and deeplearning-based variant calling have enabled long-read data to approach or, in selected genomic contexts, exceed short-read accuracy for small-variant detection [19–21]. Long reads also facilitate the joint interpretation of small and structural variants and can reduce errors caused by treating nearby variants as independent events [22, 23].

However, longer reads do not uniformly translate into more accurate small-variant calls. The relative performance of short- and long-read sequencing depends on the sequencing platform, chemistry, base-caller, coverage, aligner, variant caller, reference representation, and local genomic context [8,24]. The balance between SNV and indel accuracy is especially technology dependent. SNVs are generally well resolved by both highly accurate short reads and contemporary long-read platforms, whereas indel calling remains more sensitive to homopolymer length errors, local alignment ambiguity, repeat structure, and alternative representations of the same haplotype sequence [16, 21]. PacBio HiFi and ONT data also exhibit distinct residual error profiles, and variant callers trained for one chemistry or basecalling model may not transfer reliably to another [21]. Consequently, aggregate precision and recall values can obscure substantial context-specific differences and may provide an incomplete account of where and why a sequencing technology fails.

Benchmarking these differences is itself non-trivial. Established truth sets are most complete in well-characterized genomic regions, whereas difficult loci may contain unresolved variants, structurally divergent haplotypes, or representation-dependent discrepancies between callsets [10, 25]. Moreover, equivalent haplotypes can be encoded as different combinations of SNVs, indels, and complex variants, causing conventional comparison procedures to label biologically equivalent calls as false positives and false negatives [26, 27]. Recent haplotype-resolved assemblies, telomere-to-telomere references, pangenome resources, expanded Genome in a Bottle benchmarks, and improved variant-comparison methods have increased the accessibility of complex regions [12,23,28,29], but they have also exposed limitations in evaluation frameworks that treat small and structural variants separately or rely on exact VCF-record matching.

Previous studies have benchmarked individual small-variant callers or compared short- and long-read sequencing across selected samples and genomic stratifications [8, 24]. These analyses have demonstrated the high overall accuracy of modern pipelines and the advantages of long reads in repetitive and medically relevant regions [12,21]. Nevertheless, several questions remain insufficiently resolved. In particular, it remains unclear to what extent observed performance differences arise from read-level sequencing errors, mapping ambiguity, local sequence composition, variant representation, or caller-specific decision rules. It is also unclear whether regions conventionally classified as difficult for short reads are uniformly improved by long reads, or whether each technology exhibits distinct and partially overlapping failure modes.

Here, we systematically compare SNV and short-indel detection from representative short-read and long-read sequencing datasets using multiple alignment and variant-calling pipelines. We evaluate performance across genome-wide benchmark regions and stratify errors by sequence repetitiveness, mappability, segmental duplication, homopolymer and tandemrepeat context, local sequence complexity, variant type, and allele length. Beyond reporting aggregate precision and recall, we analyse shared and technology-specific false-positive and false-negative calls to determine how read length, sequencing-error profiles, alignment behaviour, and local genomic context shape variant-calling outcomes. By distinguishing limitations intrinsic to the sequencing data from those introduced by alignment, variant-calling, and benchmarking procedures, this study provides a mechanistic assessment of the complementary strengths and weaknesses of short- and long-read small-variant detection and identifies opportunities for improving future variant-calling methods.

## Results

### A region-stratified framework for comparing short- and long-read variant detection

To systematically investigate how genomic sequence context influences small-variant detection across sequencing technologies, we established a region-stratified benchmarking framework using short-read (SR; Illumina) and long-read (LR; PacBio HiFi) whole-genome sequencing data from the Genome in a Bottle (GIAB) reference sample HG002 (NA24385). We evaluated single-nucleotide polymorphisms (SNPs) and small insertions and deletions (INDELs) separately across multiple sequencing depths and variant callers. Reads were aligned to the GRCh38 no-alt reference genome using BWA and minimap2, respectively. Variant calling was performed using GATK, DeepVariant, and Strelka2 for SR data, and DeepVariant, Clair3, NanoCaller for LR data. Following normalization and quality filtering, callsets were evaluated against the GIAB HG002 v4.2.1 benchmark using hap.py, with precision, recall, and F1 scores calculated across genomic stratifications, including low-mappability regions (LowMap), segmental duplications (SegDup), the major histocompatibility complex (MHC), tandem repeats, and homopolymers.

To move beyond regional performance comparisons and investigate the sequence characteristics associated with differences between SR and LR variant detection, we organized the analysis into four complementary modules (Fig. 1). First, we characterized variation in detection accuracy across genomic contexts, INDEL lengths, and sequencing depths to establish the extent and distribution of performance differences between technologies and callers. Second, we investigated read-mapping ambiguity by quantifying the fraction of reads with mapping quality zero (MAPQ 0), with particular attention to variants detected by LR but missed by SR, and by examining alternative alignment locations of ambiguously mapped short reads. Third, we evaluated how the length of LowMap and SegDup tracts and the position of variants relative to tract boundaries relate to detection performance and mapping ambiguity. Finally, we examined the influence of segmental duplication architecture by relating variant recall to local paralogous sequence identity, duplication copy number, and proximity to paralogous sequence variants (PSVs). Together, these analyses provide a framework for characterizing technologydependent differences in small-variant detection and identifying the genomic features associated with reduced variant recovery in challenging regions.

**Figure 1.**
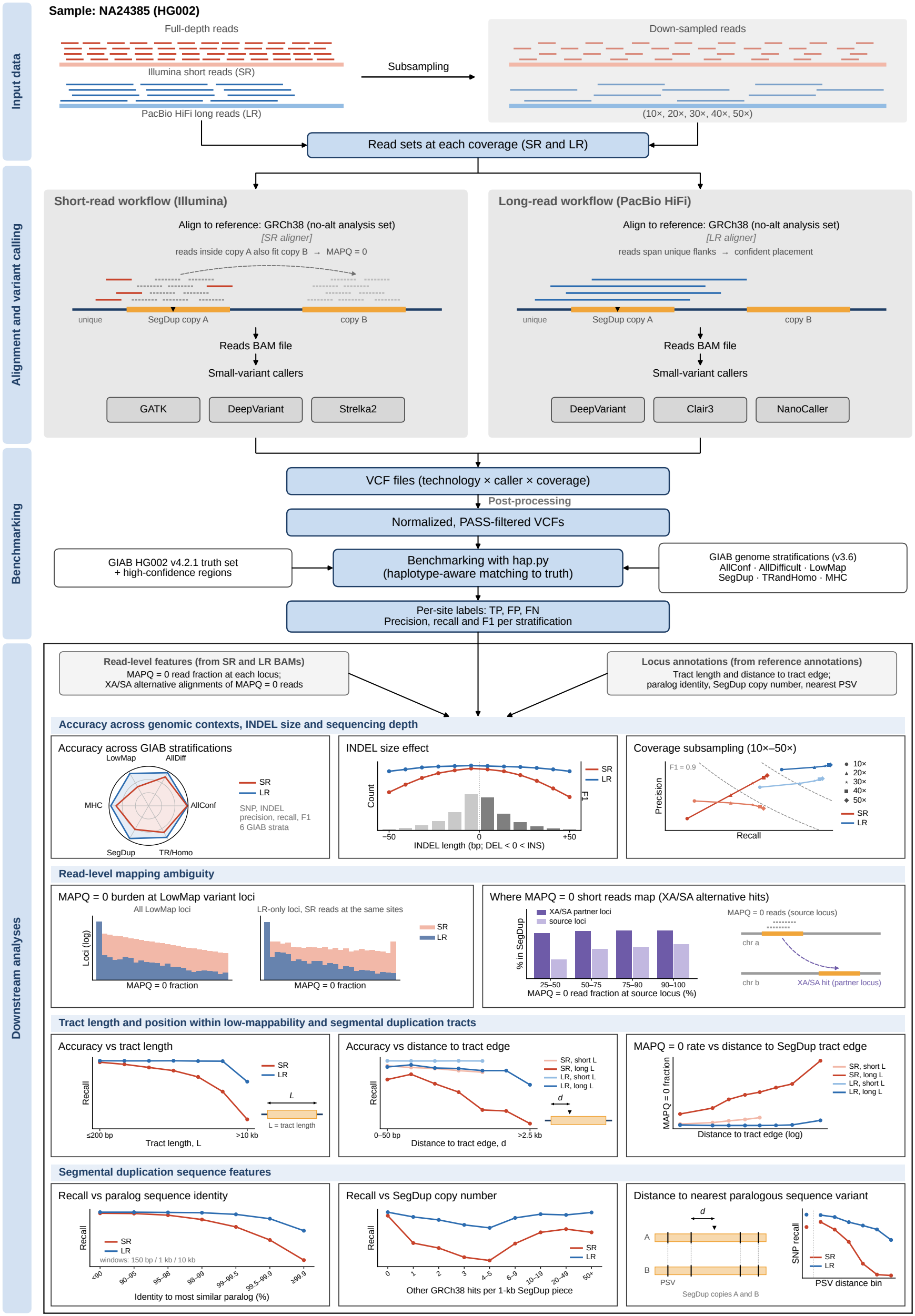
Overview of the benchmarking pipeline.

### Regional benchmarking reveals context-dependent differences between platforms and callers

We first evaluated small-variant calling performance in NA24385 across six genomic stratifications(AllConf, AllDifficult, LowMap, MHC, SegDup and TR/Homo) using hap.py-derived precision, recall, and F1 scores (Fig. 2, Table 1). Radar plots were generated separately for each sequencing technology and variant type, with F1, recall and precision shown in successive columns, allowing caller-level performance to be compared across high-confidence and difficult genomic contexts within the same sample. For SNPs, both short-read and HiFi callsets performed strongly in AllConf regions, with F1 scores extending close to the outer boundary of the radar plots(Fig. 2a,d). This indicates that, in well-characterized and largely uniquely mappable regions, Illumina-based and HiFi-based pipelines both recover small variants with high accuracy. The distinction between technologies became more pronounced in challenging regions. In LowMap and SegDup, the short-read profiles contracted inward, whereas the HiFi profiles remained closer to the outer boundary (Fig. 2a–f). This pattern was consistent across the HiFi callers tested, although individual tools differed in absolute performance. Deep-Variant and Clair3 showed particularly stable profiles across stratifications, and NanoCaller closely followed them for SNPs (Fig. 2d–f). The short-read SNP callers showed a different behavior. GATK, DeepVariant, and Strelka2 were broadly comparable in high-confidence regions, but their profiles diverged in difficult stratifications(Fig. 2a–c). However, the callers lost performance in different ways. For DeepVariant and Strelka2, the reduction was concentrated in recall, particularly in LowMap, whereas GATK retained comparatively high recall but showed a marked decline in precision in LowMap and SegDup (Fig. 2b,c). This suggests that ambiguous evidence in low-mappability sequence forces a trade-off between sensitivity and specificity that callers resolve differently: DeepVari-ant and Strelka2 tend to withhold calls at such sites, whereas GATK continues to call them at the cost of additional false positives. In contrast to LowMap and SegDup, MHC had little effect on short-read SNP calling (Fig. 2a). In practical terms, short reads appear to retain reasonable specificity when calls are made, but a larger fraction of true sites lack sufficiently reliable evidence to pass calling or genotyping thresholds. INDELs showed the same technology-level structure but with a larger overall performance penalty, consistent with the greater sensitivity of small insertion and deletion calling to local alignment quality(Fig. 2g–l). In AllConf regions, HiFi and short-read callsets again showed relatively strong performance. In LowMap and SegDup regions, however, short-read INDEL F1 and recall contracted more substantially than the corresponding HiFi metrics. TR/Homo, by contrast, had only a modest effect on short-read IN-DEL calling. The caller-specific trade-off observed for SNPs was also evident for INDELs, with DeepVariant losing mainly recall and GATK losing mainly precision in these regions(Fig. 2h,i). The HiFi INDEL profiles were not uniformly perfect across callers, but they remained more stable across difficult contexts than the short-read profiles. This was particularly evident in recall, where HiFi DeepVariant and Clair3 retained a larger fraction of truth-set INDELs in low-mappability sequence. NanoCaller performed consistently below DeepVariant and Clair3 across all stratifications, including AllConf, pointing to a caller-level offset rather than a region-specific deficit. Together, the NA24385 radar plots indicate that the long-read advantage is not confined to a single variant class. For both SNPs and INDELs, the largest separation between sequencing technologies occurred in LowMap and SegDup, where read placement is expected to be ambiguous. The stronger separation in recall than precision further suggests that short-read underperformance in NA24385 reflects unreliable evidence in difficult sequence rather than a uniform degradation of all accuracy metrics, manifesting as reduced recall or reduced precision depending on the caller. Similar patterns were observed in the evaluation of NA12878 calling results against the Platinum Pedigree truth set v1.2 benchmark (Supplementary Fig. 1, Supplementary Table 1), suggesting that the long-read advantage in difficult genomic regions is consistent across samples.

**Table 1.**
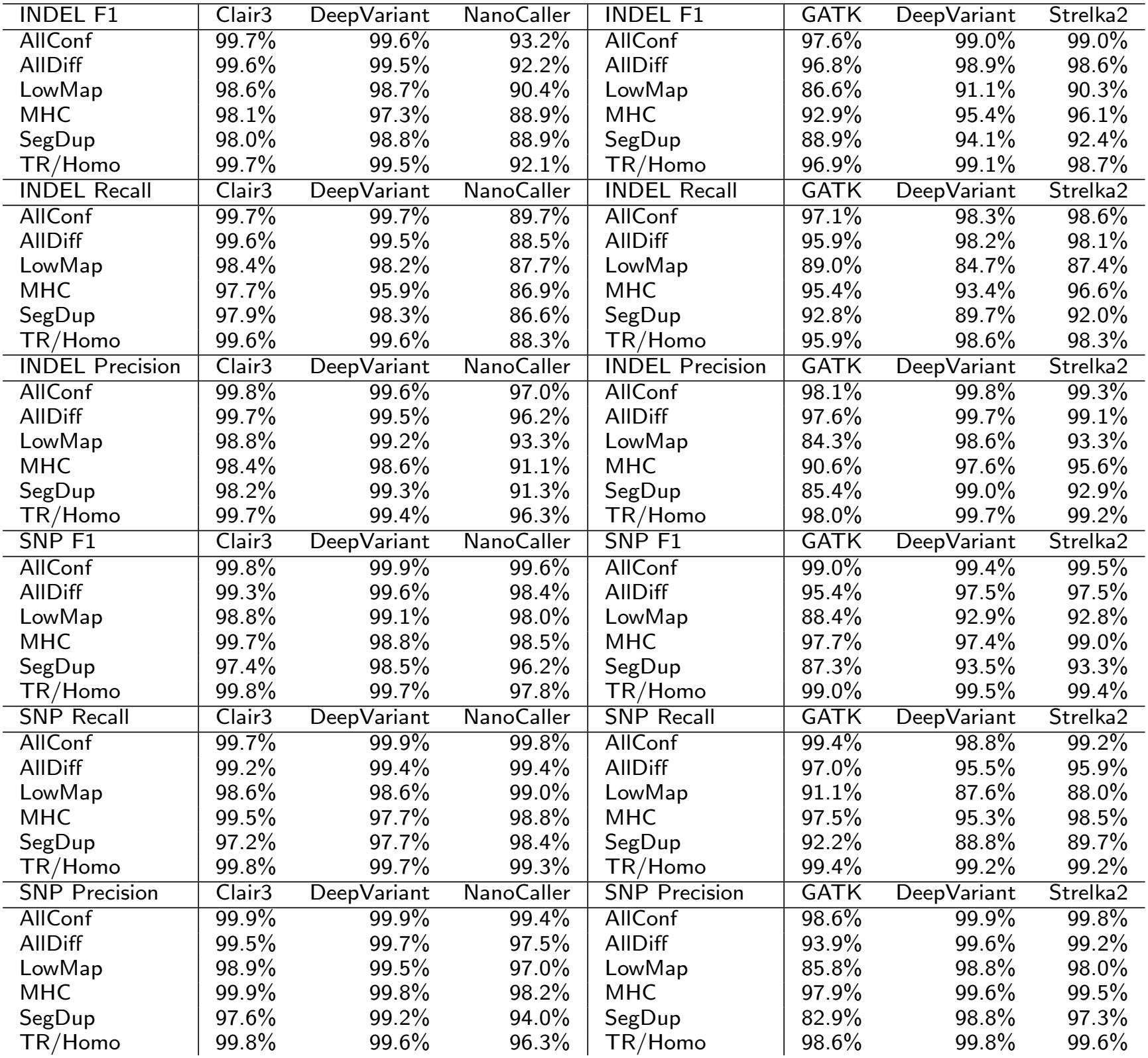
Precision, recall and F1 score of short-read (SR L1) and HiFi (HiFi L1) variant callers across genomic stratifications in NA24385, evaluated with hap.py against the GIAB HG002 v4.2.1 benchmark.

| INDEL F1 | Clair3 | DeepVariant | NanoCaller | INDEL F1 | GATK | DeepVariant | Strelka2 |
|---|---|---|---|---|---|---|---|
| AllConf | 99.7% | 99.6% | 93.2% | AllConf | 97.6% | 99.0% | 99.0% |
| AllDiff | 99.6% | 99.5% | 92.2% | AllDiff | 96.8% | 98.9% | 98.6% |
| LowMap | 98.6% | 98.7% | 90.4% | LowMap | 86.6% | 91.1% | 90.3% |
| MHC | 98.1% | 97.3% | 88.9% | MHC | 92.9% | 95.4% | 96.1% |
| SegDup | 98.0% | 98.8% | 88.9% | SegDup | 88.9% | 94.1% | 92.4% |
| TR/Homo | 99.7% | 99.5% | 92.1% | TR/Homo | 96.9% | 99.1% | 98.7% |
| INDEL Recall | Clair3 | DeepVariant | NanoCaller | INDEL Recall | GATK | DeepVariant | Strelka2 |
| AllConf | 99.7% | 99.7% | 89.7% | AllConf | 97.1% | 98.3% | 98.6% |
| AllDiff | 99.6% | 99.5% | 88.5% | AllDiff | 95.9% | 98.2% | 98.1% |
| LowMap | 98.4% | 98.2% | 87.7% | LowMap | 89.0% | 84.7% | 87.4% |
| MHC | 97.7% | 95.9% | 86.9% | MHC | 95.4% | 93.4% | 96.6% |
| SegDup | 97.9% | 98.3% | 86.6% | SegDup | 92.8% | 89.7% | 92.0% |
| TR/Homo | 99.6% | 99.6% | 88.3% | TR/Homo | 95.9% | 98.6% | 98.3% |
| INDEL Precision | Clair3 | DeepVariant | NanoCaller | INDEL Precision | GATK | DeepVariant | Strelka2 |
| AllConf | 99.8% | 99.6% | 97.0% | AllConf | 98.1% | 99.8% | 99.3% |
| AllDiff | 99.7% | 99.5% | 96.2% | AllDiff | 97.6% | 99.7% | 99.1% |
| LowMap | 98.8% | 99.2% | 93.3% | LowMap | 84.3% | 98.6% | 93.3% |
| MHC | 98.4% | 98.6% | 91.1% | MHC | 90.6% | 97.6% | 95.6% |
| SegDup | 98.2% | 99.3% | 91.3% | SegDup | 85.4% | 99.0% | 92.9% |
| TR/Homo | 99.7% | 99.4% | 96.3% | TR/Homo | 98.0% | 99.7% | 99.2% |
| SNP F1 | Clair3 | DeepVariant | NanoCaller | SNP F1 | GATK | DeepVariant | Strelka2 |
| AllConf | 99.8% | 99.9% | 99.6% | AllConf | 99.0% | 99.4% | 99.5% |
| AllDiff | 99.3% | 99.6% | 98.4% | AllDiff | 95.4% | 97.5% | 97.5% |
| LowMap | 98.8% | 99.1% | 98.0% | LowMap | 88.4% | 92.9% | 92.8% |
| MHC | 99.7% | 98.8% | 98.5% | MHC | 97.7% | 97.4% | 99.0% |
| SegDup | 97.4% | 98.5% | 96.2% | SegDup | 87.3% | 93.5% | 93.3% |
| TR/Homo | 99.8% | 99.7% | 97.8% | TR/Homo | 99.0% | 99.5% | 99.4% |
| SNP Recall | Clair3 | DeepVariant | NanoCaller | SNP Recall | GATK | DeepVariant | Strelka2 |
| AllConf | 99.7% | 99.9% | 99.8% | AllConf | 99.4% | 98.8% | 99.2% |
| AllDiff | 99.2% | 99.4% | 99.4% | AllDiff | 97.0% | 95.5% | 95.9% |
| LowMap | 98.6% | 98.6% | 99.0% | LowMap | 91.1% | 87.6% | 88.0% |
| MHC | 99.5% | 97.7% | 98.8% | MHC | 97.5% | 95.3% | 98.5% |
| SegDup | 97.2% | 97.7% | 98.4% | SegDup | 92.2% | 88.8% | 89.7% |
| TR/Homo | 99.8% | 99.7% | 99.3% | TR/Homo | 99.4% | 99.2% | 99.2% |
| SNP Precision | Clair3 | DeepVariant | NanoCaller | SNP Precision | GATK | DeepVariant | Strelka2 |
| AllConf | 99.9% | 99.9% | 99.4% | AllConf | 98.6% | 99.9% | 99.8% |
| AllDiff | 99.5% | 99.7% | 97.5% | AllDiff | 93.9% | 99.6% | 99.2% |
| LowMap | 98.9% | 99.5% | 97.0% | LowMap | 85.8% | 98.8% | 98.0% |
| MHC | 99.9% | 99.8% | 98.2% | MHC | 97.9% | 99.6% | 99.5% |
| SegDup | 97.6% | 99.2% | 94.0% | SegDup | 82.9% | 98.8% | 97.3% |
| TR/Homo | 99.8% | 99.6% | 96.3% | TR/Homo | 98.6% | 99.8% | 99.6% |

**Figure 2.**
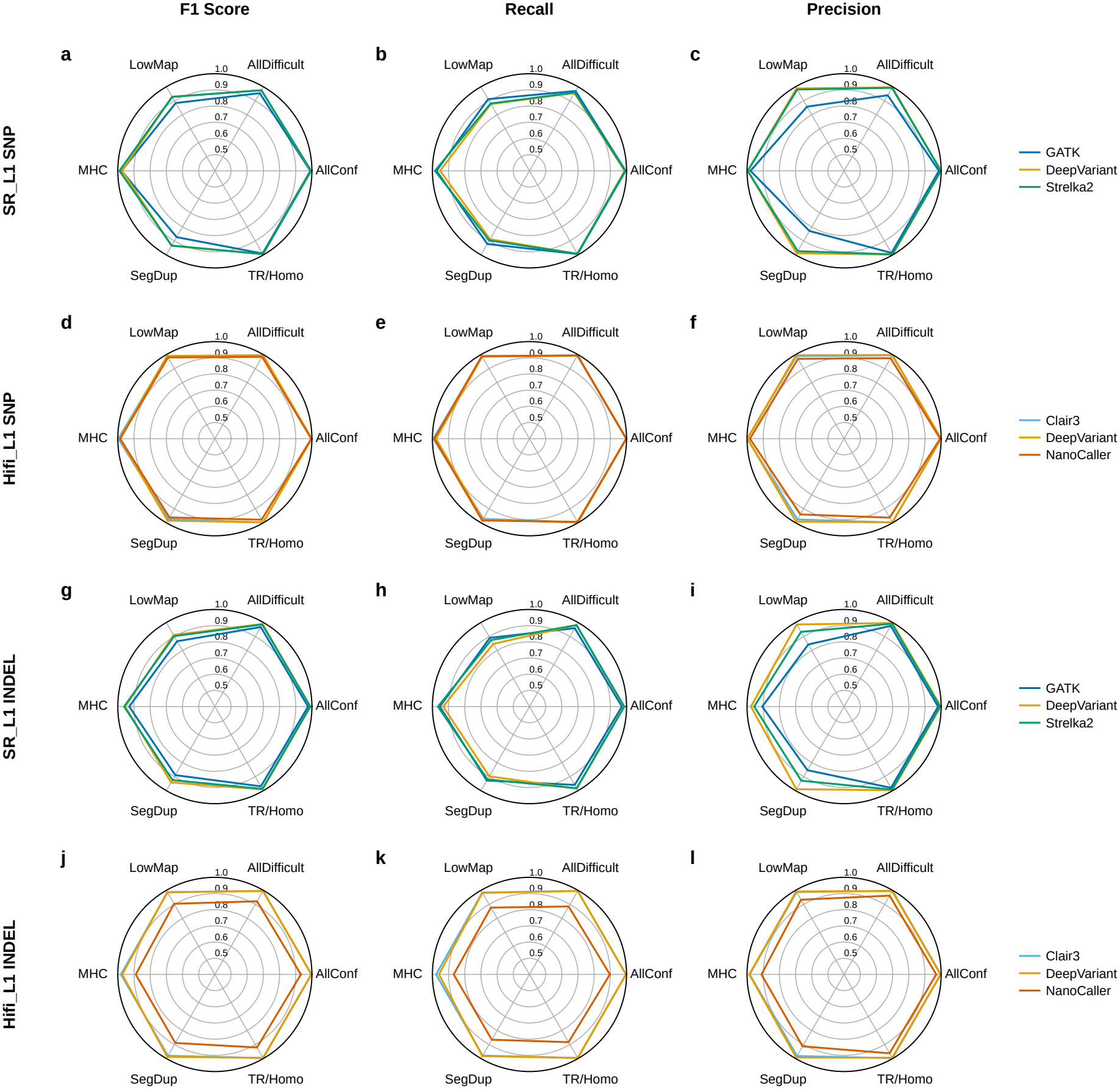
Small-variant calling performance across genomic stratifications. Radar plots of F1 score (left column), recall (middle column) and precision (right column) for the short-read (SR L1; Illumina) and long-read (HiFi L1; PacBio HiFi) datasets of NA24385, evaluated with hap.py against the GIAB HG002 v4.2.1 benchmark in six stratifications: AllConf, AllDifficult, LowMap, MHC, SegDup and TR/Homo. Concentric circles mark values from 0.5 to 1.0. **a–c** SR SNPs called with GATK, DeepVariant and Strelka2. **d–f** HiFi SNPs called with Clair3, DeepVariant and NanoCaller. **g–i** SR INDELs called with GATK, DeepVariant and Strelka2. **j–l** HiFi INDELs called with Clair3, DeepVariant and NanoCaller.

### Sequencing depth and INDEL length reveal further platform- and caller-specific differences

To assess the dependence of variant detection on sequencing depth, we compared SNP calling performance across subsampled datasets spanning 10× to 50× coverage, separately for AllConf and AllDifficult regions (Fig. 3a–d). Increasing depth generally improved detection, but the magnitude and balance of gains in precision and recall varied between platforms and callers. HiFi-based DeepVariant and Clair3 already achieved high precision and recall at the lowest evaluated depth, with comparatively modest gains as coverage increased. Their trajectories clustered at higher depths, consistent with diminishing returns over the evaluated range.

**Figure 3.**
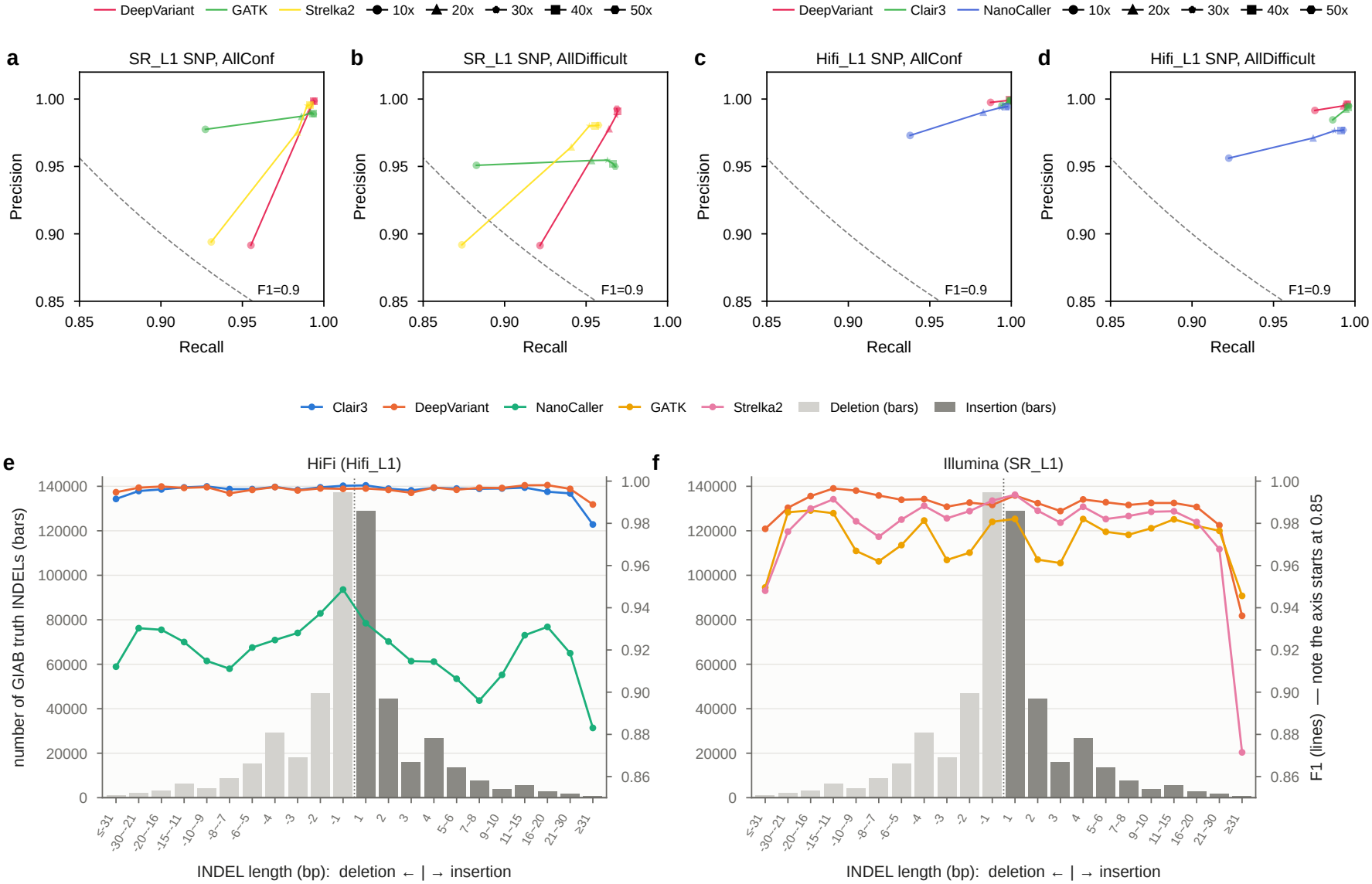
Effects of sequencing depth and INDEL length on calling performance. **a–d** Precision against recall for SNPs called from SR L1 (**a**,**b**) and HiFi L1 (**c**,**d**) after subsampling to 10×, 20×, 30×, 40× and 50× coverage, in AllConf (**a**,**c**) and AllDifficult (**b**,**d**). Lines connect the depths of each caller, and marker shapes indicate depth. Dashed curve, F1 = 0.9. **e**,**f** F1 score (lines, right axis) of INDELs called from HiFi L1 (**e**) and SR L1 (**f**) in AllConf, by INDEL length. Negative lengths denote deletions and positive lengths insertions. Bars (left axis) show the number of GIAB benchmark INDELs in each length bin (light grey, deletions; dark grey, insertions). The right axis starts at 0.85.

SR-based DeepVariant and Strelka2 showed larger improvements from low to higher coverage, reflecting a stronger dependence on depth, and in AllConf regions converged with the other callers at higher depths. This response was not uniform across methods: GATK gained recall with increasing depth without a corresponding sustained increase in precision, while HiFi-based NanoCaller remained below DeepVariant and Clair3 despite improvement with additional coverage. In AllDifficult regions, however, additional depth did not close the gap between platforms: short-read callers improved with coverage but plateaued at lower recall, even at 50× (Fig. 3b). GATK, in particular, gained recall with increasing depth without a corresponding increase in precision. By contrast, HiFi DeepVariant achieved higher recall at 10× than any short-read caller at 50× (Fig. 3b,d). These results show that the benefit of additional sequencing depends on the platform and caller, and that increasing depth does not eliminate the short-read deficit in difficult regions, suggesting that additional short reads in these regions contribute little usable evidence, a possibility we examine below.

Beyond genomic context, we examined how INDEL length affects calling performance (Fig. 3e,f). The GIAB truth set is dominated by 1–2-bp INDELs, and the number of truth INDELs declines sharply beyond 10 bp, such that aggregate INDEL metrics largely reflect performance on short events. HiFi DeepVariant and Clair3 maintained near-uniform F1 across deletions and insertions up to 30 bp, declining only for the longest insertions. NanoCaller remained below these callers across all length bins without a consistent length dependence, in line with the caller-level offset observed in Fig. 2. Among short-read callers, DeepVariant was the most accurate and stable across short and intermediate lengths, whereas GATK and Strelka2 fluctuated more in the 2–10-bp range, possibly reflecting sensitivity to length changes in tandem repeats and homopolymers. All short-read callers declined markedly for INDELs longer than 30 bp, with a stronger reduction for insertions than for deletions. This asymmetry is consistent with previous reports that short-read sensitivity declines with increasing insertion size, likely because reads spanning long insertions align less efficiently than those spanning deletions [24]. Thus, HiFi INDEL detection was largely insensitive to length, whereas short-read detection varied at intermediate lengths and deteriorated for long insertions, differences that are largely masked in aggregate metrics because such events are comparatively rare.

### Short-read mapping ambiguity accompanies a subset of variants recovered only by long reads

To test whether the stratified performance gap in NA24385 was associated with read-level alignment ambiguity, we quantified the fraction of overlapping reads with mapping quality equal to zero at low-mappability variant loci. MAPQ=0 fraction was used as a direct proxy for ambiguous read placement, with higher values indicating that a larger proportion of reads at a locus could not be confidently assigned to a unique genomic position. We compared these distributions at all low-mappability variant loci and at loci called by only one technology, separately for SNPs and INDELs. Across all low-mappability variant loci, the HiFi and short-read distributions were clearly separated (Fig. 4a,b). For SNPs, the HiFi analysis included 190,930 loci, while the short-read analysis included 170,930 loci. For INDELs, the corresponding counts were 9,908 HiFi loci and 8,676 short-read loci. HiFi loci were strongly enriched near MAPQ=0 fraction of zero, indicating that most variant-supporting long reads were placed with high mapping confidence even within low-mappability regions. In contrast, the short-read distributions extended across the full range of MAPQ=0 fractions, with substantial signal persisting at high values, including fractions above 0.6 and 0.8. This broad right-shift in the short-read distribution provides a read-level explanation for the reduced short-read performance, especially the reduced recall observed in the radar plots. A variant locus can have nominal read coverage but still provide limited usable evidence if many overlapping reads have ambiguous placement. In such cases, additional reads do not necessarily translate into informative allele support. Instead, they increase the pool of low-confidence alignments, reducing the effective depth available for accurate genotyping, consistent with the limited benefit of additional short-read coverage in difficult regions (Fig. 3b). The HiFi distribution shows the opposite behavior: most low-mappability loci retain a low MAPQ=0 burden, consistent with long reads spanning enough flanking sequence to resolve placement across repetitive or duplicated regions. We then focused specifically on LR-only loci, defined as positions present in the long-read variant set but absent from the short-read variant set, and extracted shortread MAPQ=0 fractions from the BAM at the same genomic coordinates (Fig. 4c,d). At these loci, long reads were almost uniformly placed with high confidence, with a median MAPQ=0 fraction of zero, whereas short reads at the same SNP positions had a median of 0.082. Short reads overlapping the same positions showed a much broader distribution across the full MAPQ=0 fraction range, including a distinct group of loci at which essentially all overlapping short reads had MAPQ=0. This comparison is especially informative because it holds genomic position constant. The observed difference therefore cannot be attributed solely to different sets of regions being sampled by the two technologies. Instead, it indicates that at the same low-mappability coordinates where long reads provide confident support for a variant, short reads frequently remain ambiguously aligned.The INDEL analysis showed the same qualitative pattern, with smaller absolute counts reflecting the lower number of INDEL loci. HiFi-supported INDEL positions were again enriched for low MAPQ=0 fractions, whereas short reads at the corresponding or broader low-mappability INDEL loci showed a more diffuse distribution extending toward high MAPQ=0 values. This reinforces the interpretation that ambiguous read placement affects both SNP and INDEL detection in NA24385, although the impact is particularly consequential for INDELs because local alignment uncertainty directly affects breakpoint and allele representation. We also examined SR-only loci, defined as positions present in the short-read variant set but absent from the long-read variant set (Fig. 4e,f). At these loci, the pattern was reversed: short reads showed a low MAPQ=0 burden, whereas long reads displayed a broad tail that exceeded the short-read distribution at intermediate and high MAPQ=0 fractions, for both SNPs and INDELs. This suggests that some SR-only calls lie in sequence where even long reads cannot be placed uniquely, consistent with the decline in HiFi performance within the longest and most similar duplications described below. Because these loci were defined from callsets, they may also include short-read false positives arising from confidently but incorrectly placed reads, and the two explanations are not distinguished here. Overall, the MAPQ=0 analysis links the NA24385 stratified bench-marking results to a concrete alignment-level mechanism. The radar plots show that short-read callsets lose performance primarily in difficult regions, especially, through reduced recall or reduced precision depending on the caller. The MAPQ=0 distributions show that these same regions contain a substantially higher burden of ambiguously mapped short reads. Long reads, by contrast, retain confident placement at most low-mappability variant loci, providing a mechanistic basis for their more complete variant recovery in NA24385, although the SR-only loci indicate that this advantage does not extend to all low-mappability sequence.

**Figure 4.**
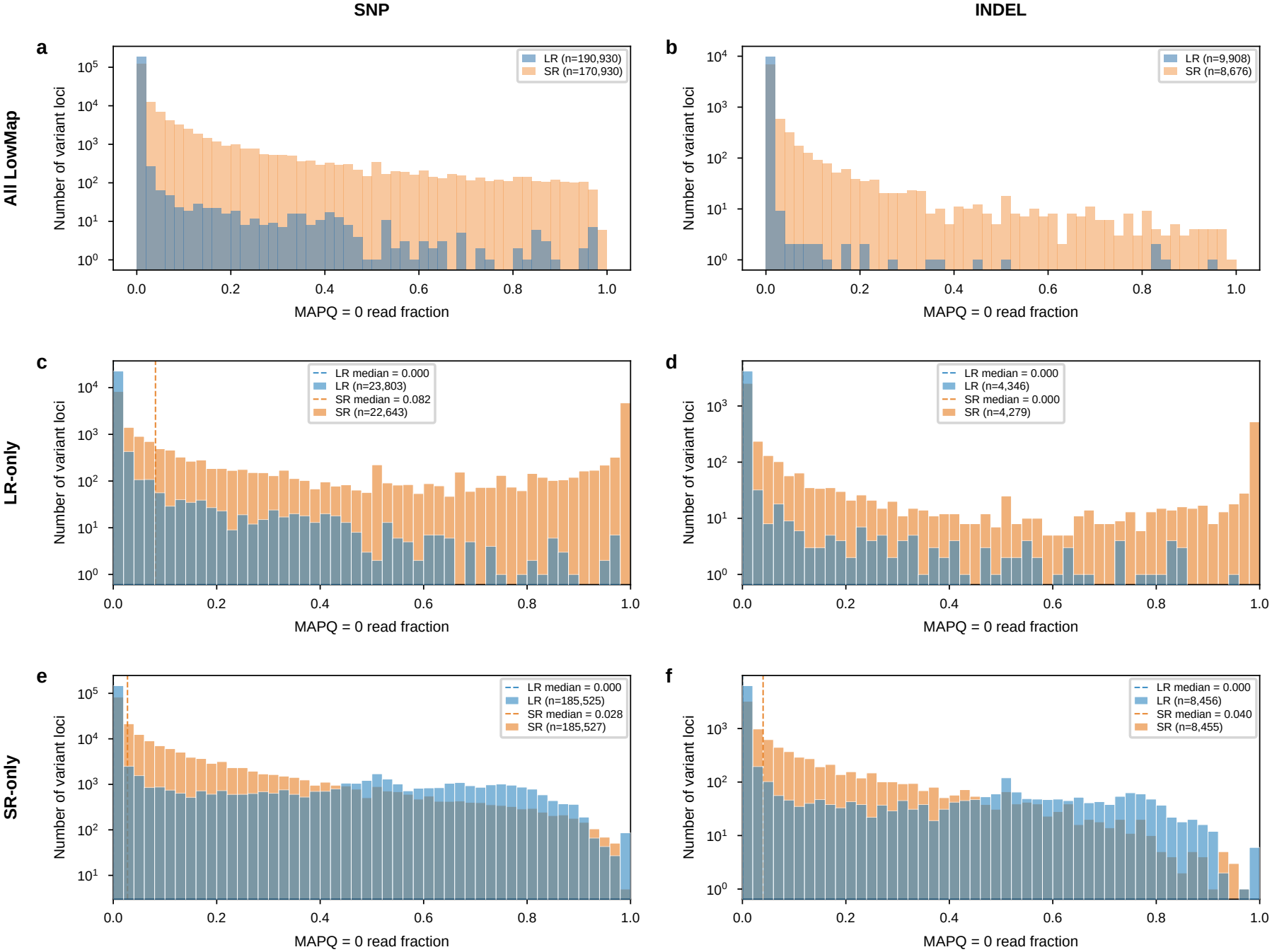
Mapping ambiguity of short and long reads at low-mappability variant loci. Distributions of the fraction of overlapping reads with MAPQ 0 at variant loci in LowMap, for SNPs (left column) and INDELs (right column), computed from the SR (orange) and LR (blue) alignments of NA24385. Variant loci were taken from the DeepVariant callsets of each platform. The vertical axis is on a log scale, and *n* is the number of loci. **a**,**b** All variant loci called from each platform. **c**,**d** LR-only loci, called from LR but not SR data, with the MAPQ 0 fraction computed from the LR and SR alignments at the same positions. **e**,**f** SR-only loci, called from SR but not LR data, computed in the same way. Dashed lines, medians.

### Short-read recall declines with increasing alignment ambiguity in long tract interiors

Using the HG002 SR and HiFi DeepVariant callsets, we examined how variant-calling performance within benchmark high-confidence regions varied with tract length and distance to the nearest tract edge in LowMap and SegDup (Methods). SR recall generally decreased with increasing tract length, most markedly in LowMap, where recall for both SNPs and INDELs fell below 50% in tracts longer than 10 kb, compared with approximately 80% for HiFi (Supplementary Fig. 2). Precision varied less with tract length, and the reductions in F1 largely followed those in recall. Differences between technologies were smaller in SegDup.

Within tract-length groups, SR recall in LowMap generally declined with distance from the nearest tract edge, with the largest losses in tracts longer than 10 kb (Fig. 5b,d). In these tracts, SR SNP recall decreased from above 80% near the edge to approximately 40% at distances greater than 2.5 kb. INDEL recall was also below 50% in the most distant evaluable ranges, although fewer ranges met the minimum denominator. HiFi recall remained close to 100% in tracts up to 10 kb and decreased mainly in the interiors of the longest LowMap tracts. Precision changed less than recall with distance from the tract edge (Fig. 5a,c). At distances greater than 2.5 kb in tracts longer than 10 kb, SR precision remained above 80% for both SNPs and INDELs while recall fell below 50%, suggesting that the reduced performance mainly reflected missed benchmark variants.

**Figure 5.**
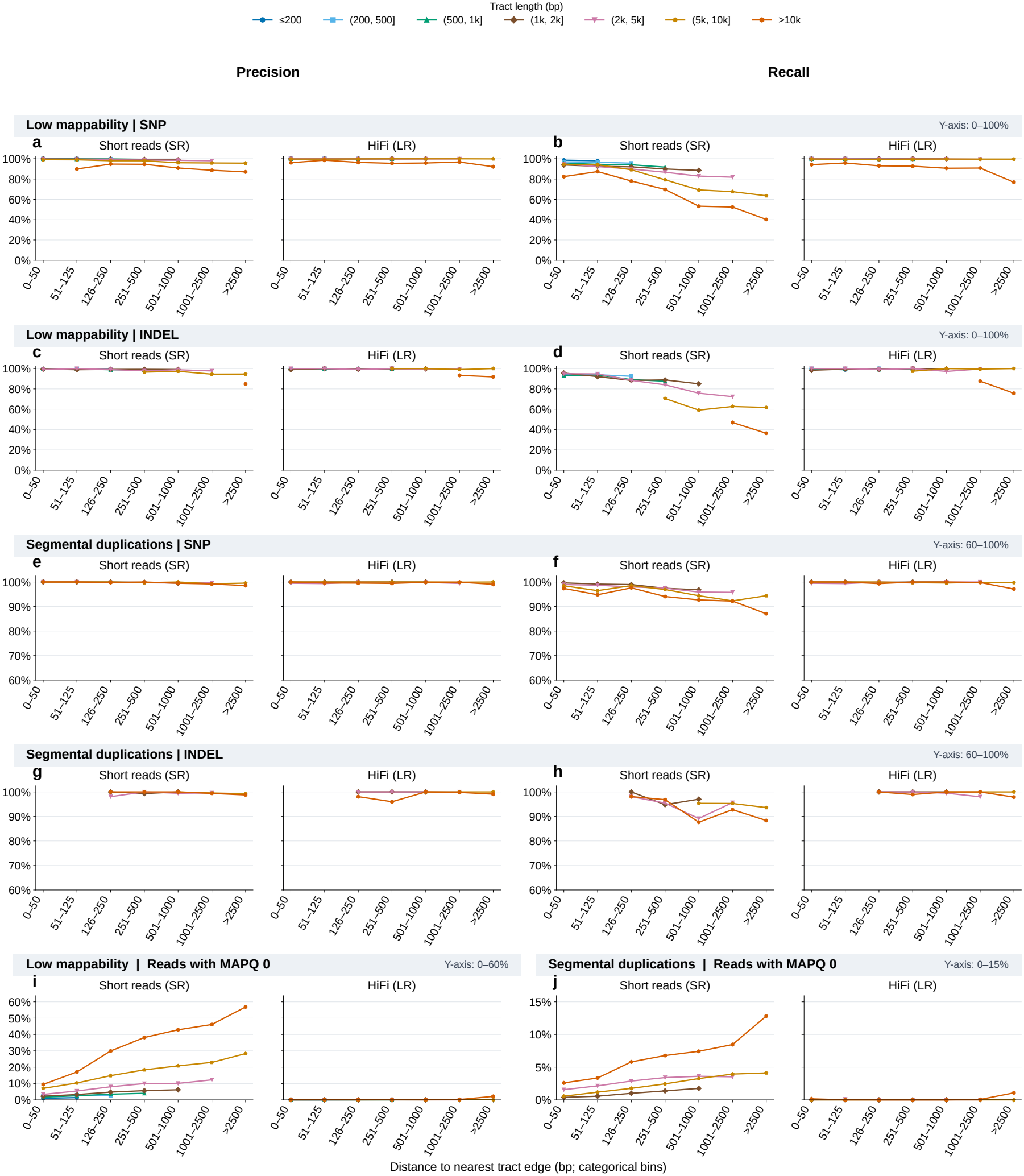
Distance to the tract edge in relation to variant recovery and alignment ambiguity in LowMap and SegDup. **a–h** Precision (**a, c, e, g**) and recall (**b, d, f, h**) of the SR and LR DeepVariant callsets for SNPs (**a, b, e, f**) and INDELs (**c, d, g, h**) in LowMap (**a–d**) and SegDup (**e–h**), with variants grouped by distance to the nearest tract edge and by tract length (colors). Values are shown where the denominator contained at least 50 records. **i**,**j** Percentage of SR and LR reads with MAPQ 0 in LowMap (**i**) and SegDup (**j**), grouped by distance from the read midpoint to the nearest tract edge and by tract length. Combinations with fewer than 100 reads are not shown. Segmental duplications are at least 1 kb long, so only tract-length groups above 1 kb occur in **e–h** and **j**. Y-axis ranges differ between rows and are given at the right of each row header.

In SegDup, distance-associated changes were smaller (Fig. 5e–h). SR SNP recall generally declined toward tract interiors and was lowest in the interiors of tracts longer than 10 kb, whereas HiFi recall remained near 100% in most distance ranges. SR INDEL recall varied more across distance ranges, and precision remained high for both technologies.

We next examined whether alignment ambiguity varied with position within LowMap and SegDup tracts, using the fraction of reads with MAPQ 0 in the SR and LR alignments, grouped by distance to the nearest tract edge and by tract length (Fig. 5i,j; Methods). For SR, the MAPQ 0 fraction generally increased with distance from the tract edge in both region types, most markedly in the longest tracts. In SegDup tracts longer than 10 kb, the fraction rose from 2.6% near the edge to 12.8% at distances greater than 2.5 kb (Fig. 5j). The increase was larger in LowMap, where the fraction in tracts longer than 10 kb reached 57% in the tract interior (Fig. 5i). At each distance, tracts longer than 10 kb had higher MAPQ 0 fractions than shorter tracts; in LowMap tracts of 1 kb or less, the fraction remained below 5%.

In contrast, LR alignments changed much less with tract position or length. In both LowMap and SegDup, the MAPQ 0 fraction for HiFi reads remained close to zero in most distance ranges, with small increases at distances greater than 2.5 kb in tracts longer than 10 kb (approximately 2% in LowMap and 1% in SegDup). The decline in SR recall toward the interiors of long tracts thus coincided with greater shortread alignment ambiguity, whereas HiFi alignments retained low MAPQ 0 fractions and higher recall.

### Sequence similarity further resolves detection difficulty within segmental duplications

We further characterized segmental duplications by the number of alternative genomic matches and by similarity to a competing sequence (Fig. 6). Grouping 1-kb duplication fragments by the number of other positions to which they aligned in GRCh38 revealed a non-monotonic relationship with recall (Fig. 6a). Fragments with no other alignment in GRCh38 were recovered almost completely by both platforms. Both platforms showed declining recall across the lower multiplicity groups, reaching a minimum in the group with four to five alternative matches, followed by recovery in groups with more matches. HiFi retained higher recall throughout these groups, whereas precision differed only modestly between platforms and did not show a uniform LR advantage (Fig. 6b). Alternative-match count alone therefore did not order duplicated regions along a simple gradient of detection difficulty. Sequence similarity showed a clearer relationship with variant recovery. Across 150-bp, 1-kb and 10-kb windows, recall declined as identity to the most confusable partner increased (Fig. 6c–e). Illumina showed larger losses than HiFi, while HiFi also deteriorated in the highest-identity groups. The decline was particularly pronounced when high similarity extended across the 10-kb window, a span comparable to HiFi read length, demonstrating that even LR detection was challenged in regions with highly similar sequence over an extended span.

**Figure 6.**
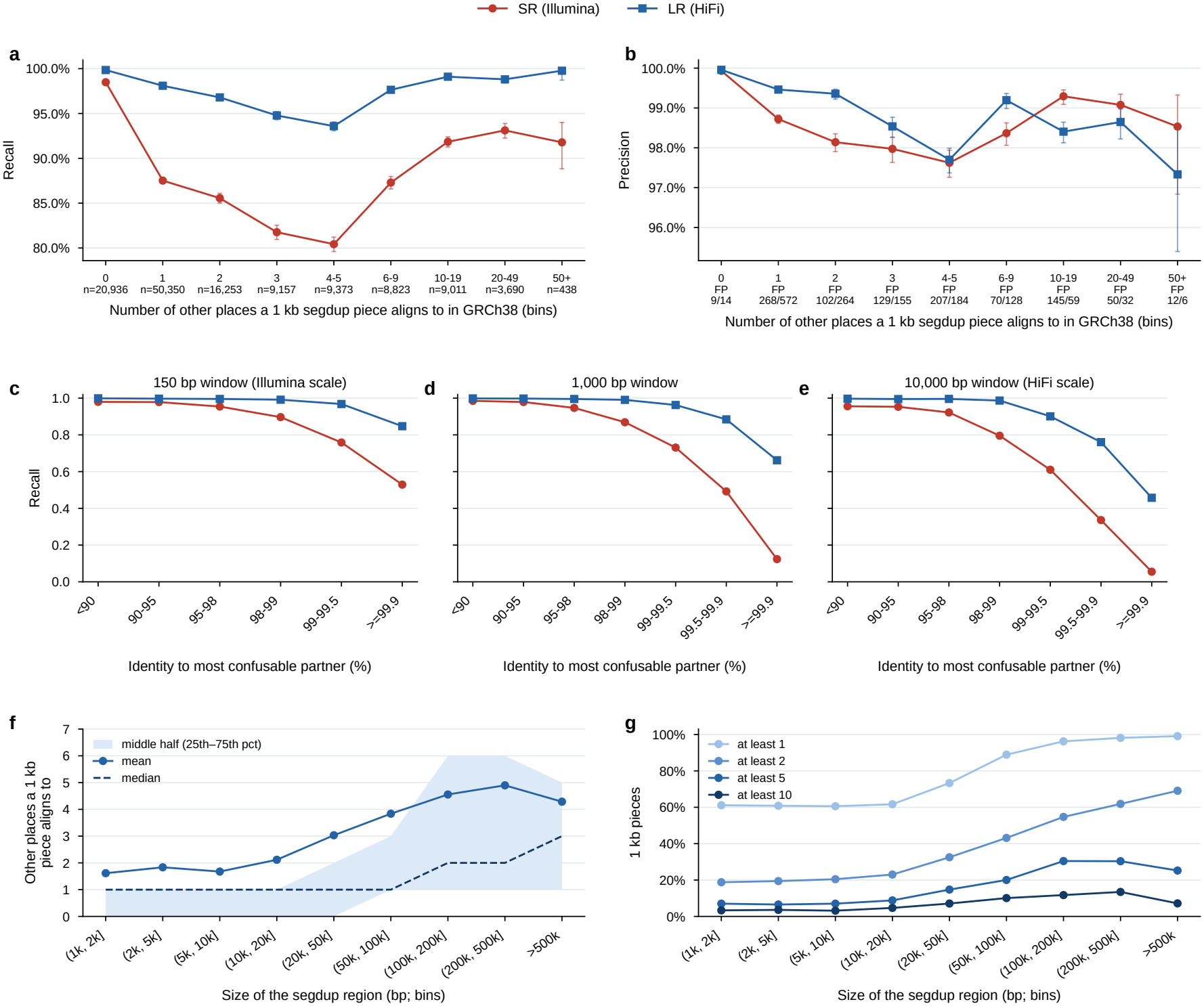
Copy number and paralog identity of segmental duplications in relation to variant recovery. **a**,**b** Recall (**a**) and precision (**b**) of the SR and LR DeepVariant callsets in SegDup, with variants grouped by the number of other GRCh38 locations to which the 1-kb SegDup window containing them aligns. In **a**, *n* is the number of benchmark variants in each group. In **b**, the number of false-positive calls is shown for SR/LR. Error bars, 95% Wilson confidence intervals. **c–e** Recall of SR and LR calls by sequence identity between the window containing each variant and its closest paralog, for windows of 150 bp (**c**), 1 kb (**d**) and 10 kb (**e**). **f** Number of other GRCh38 locations to which each 1-kb window aligns, by the size of the SegDup interval containing it. Line, mean. Dashed line, median. Shading, interquartile range. **g** Percentage of 1-kb windows that align to at least 1, 2, 5 and 10 other locations, by SegDup interval size.

We then related duplication multiplicity to the size of the segmental duplication region (Fig. 6f,g). In smaller regions, a typical 1-kb fragment aligned to only one other location, and in regions shorter than approximately 20 kb a substantial fraction of fragments had no other match in GRCh38. Multiplicity increased with region size beyond this range, with both the mean number of alternative matches and the proportion of fragments aligning to at least two other locations rising in the largest regions. Fragments with ten or more alternative matches, however, remained uncommon across all size classes. Fragments without alternative matches, which were recovered almost completely in Fig. 6a, were thus largely absent from the largest regions, whereas the longest tracts, in which short-read recall was lowest in the boundary analysis, were enriched for multi-copy sequence. Tract length therefore partly co-varies with multiplicity, although neither feature alone captured the sequence distinguishability that most closely tracked recall.

These observations refine the regional comparisons by showing that duplicated regions differ in their effects on variant detection according to both spatial context and sequence similarity. HiFi preserved greater variant recovery across these contexts, but its absolute performance declined where long stretches of sequence offered little differentiation from competing genomic locations. Together with the boundary analysis, these results support the availability of distinguishing sequence context as an organizing explanation for the observed SR–LR performance differences.

### Distance to paralogous sequence variants is associated with variant recovery within segmental duplications

Within a segmental duplication, paralogous copies can be distinguished only at positions where their sequences differ. These positions, paralogous sequence variants (PSVs), are properties of the reference genome and are distinct from the sample variants evaluated against the benchmark (Fig. 7a). A read that overlaps a PSV carries a base consistent with only one copy, whereas a read that lies entirely between PSVs matches each copy equally well. We therefore examined whether variant detection in segmental duplications varied with the distance between a variant and the nearest PSV.

**Figure 7.**
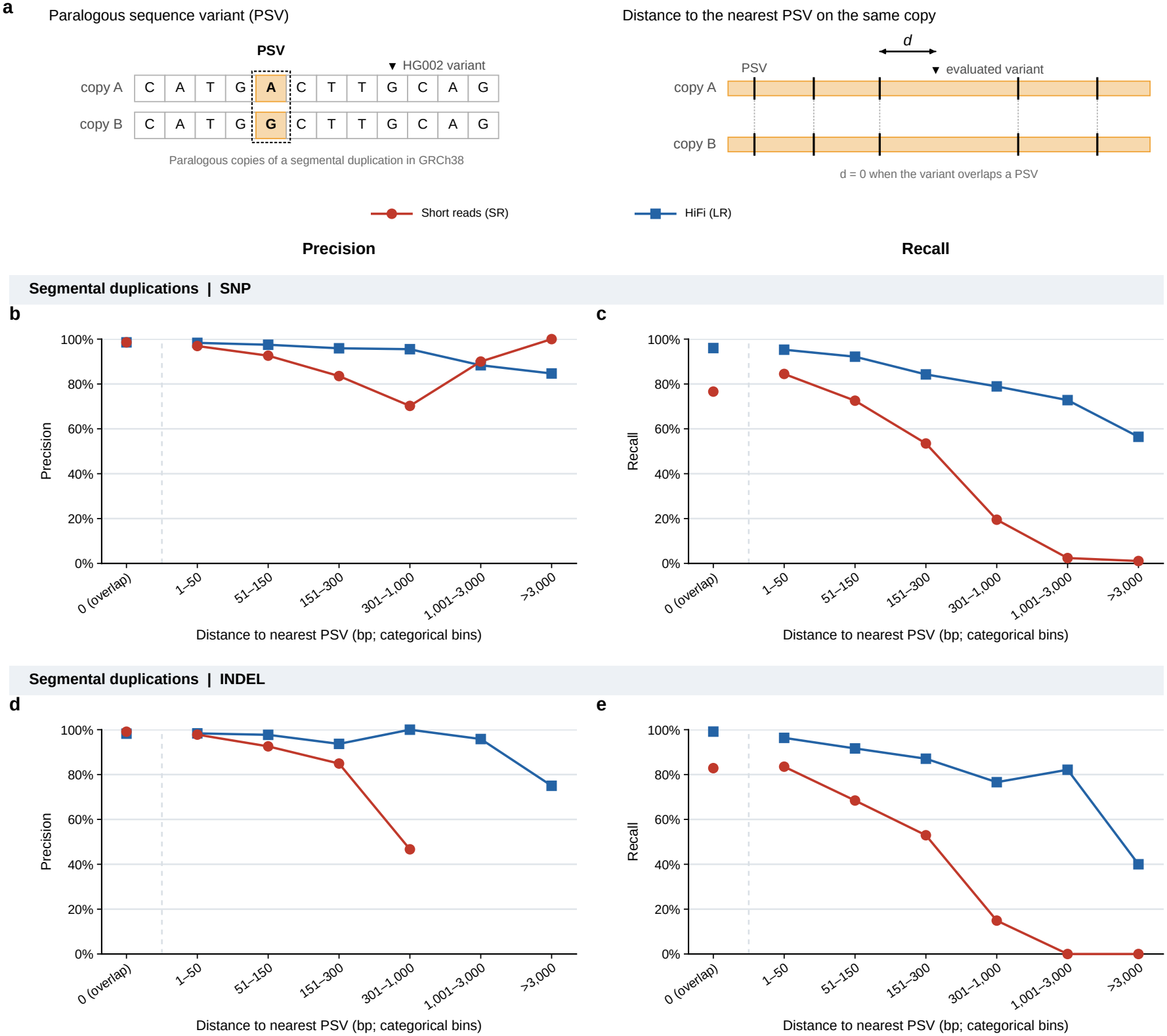
Distance to the nearest paralogous sequence variant in relation to variant recovery in segmental duplications. **a** Left, a paralogous sequence variant (PSV) is a position at which paralogous copies of a segmental duplication differ in GRCh38; it is distinct from a variant of the sample (HG002) relative to the reference. Right, for each evaluated variant, the distance *d* to the nearest PSV was measured on the same duplicated copy; *d* = 0 when the variant overlaps a PSV. **b–e** Precision (**b, d**) and recall (**c, e**) of the SR and LR DeepVariant callsets in SegDup for SNPs (**b, c**) and INDELs (**d, e**), with variants grouped by *d*. Variants overlapping a PSV (0 bp) are shown separately from the positive-distance ranges. Recall was calculated from benchmark variants and precision from called variants, within SegDup in the GIAB benchmark region covered by the PSV catalogue (38.9 Mb). The gap in **d** indicates distance ranges without SR INDEL calls.

PSVs were taken from the reference-derived catalogue released with DuploMap [30], so the same PSVs were used for both platforms. For the SR and HiFi DeepVariant callsets, we measured the distance from each benchmark variant and each called variant to the nearest PSV on the same duplicated copy, within segmental duplications in the GIAB benchmark region covered by the catalogue (38.9 Mb), and calculated recall and precision within distance ranges (Methods). Most benchmark variants lay close to a PSV: approximately 84% of SNPs and 87% of INDELs overlapped a PSV or lay within 50 bp of one, and about 1% lay more than 1 kb away.

SR recall decreased with distance for both SNPs and INDELs (Fig. 7c,e), with the largest decrease between the 151–300 and 301–1,000 bp ranges. Beyond 1 kb, SR recovered 11 of 561 SNPs and none of 43 INDELs. HiFi recall was higher than SR recall in every range, including PSV-overlapping sites, and decreased more gradually. Precision varied less with distance (Fig. 7b,d). Within 50 bp of a PSV, precision exceeded 96% for both platforms; from 50 bp to 1 kb, SR precision decreased, whereas HiFi precision remained above 93%. Beyond 1 kb, SR made too few calls for precision to be estimated reliably.

Overall, variant recovery in segmental duplications was associated with distance to the nearest PSV, and the association was stronger for SR than for HiFi. The largest decrease in SR recall occurred at distances comparable to or greater than the median short-read fragment length (404 bp; Methods), where a read pair that overlaps a variant is less likely to also overlap a PSV. This pattern is consistent with a dependence of shortread placement on copy-distinguishing bases, although the PSVs were derived from the reference rather than verified in HG002.

## Discussion

In this study, we compared SR and LR small-variant calling in HG002 across genomic contexts and examined the read-level and sequence-level features associated with the differences between the two platforms. In high-confidence regions, both platforms achieved high accuracy, and the differences were concentrated in low-mappability regions and segmental duplications (Fig. 2). Within these regions, variants detected only from LR data were frequently located at positions where SR reads had a MAPQ of 0 (Fig. 4), and SR recall declined toward the interior of long LowMap and SegDup intervals, where the proportion of MAPQ 0 reads also increased (Fig. 5). In SegDup, SR recall decreased sharply as sequence identity to the closest paralog increased, whereas LR recall remained high until identity approached 100% (Fig. 6). Together, these results indicate that the SR deficit in difficult regions arises largely from ambiguous read placement, and that its size can be anticipated from the length and paralog similarity of the surrounding sequence.

The dependence of SR recall on paralog identity can be explained by the number of paralogous sequence variants (PSVs), the positions at which duplicated copies differ, that a single read spans. An aligner can place a read in the correct copy only when the read covers at least one PSV, and Prodanov and Bansal showed that explicitly using PSVs improves long-read alignment and variant calling in segmental duplications [30]. At 99.9% identity, copies differ on average at one position per kilobase, so most 250-bp reads contain no PSV, whereas a HiFi read of 10–20 kb is expected to contain several. This is consistent with the gap between platforms widening in the highest identity ranges (Fig. 6c–e), and with SR reads near the ends of a duplication, which can extend into unique flanking sequence, being placed more reliably than reads in its interior (Fig. 5). Copy number showed a less direct relationship: SR recall was lowest for segments with four to five alternative matches and partly recovered at higher copy numbers (Fig. 6a), suggesting that the number of copies matters less than how similar they are. We did not, however, test this relationship directly.

These findings refine the general view that long reads are advantageous in repetitive regions [8, 21]. In our data, the LR advantage was not uniform across difficult regions but was concentrated in LowMap and in segmental duplications with high paralog identity, whereas the differences in tandem repeats and the MHC were smaller for most callers (Fig. 2). Outside these contexts, SR calling with current callers at 30 *×* or higher coverage approached the accuracy of LR calling (Fig. 3a–d). SR calling was also less accurate for insertions longer than 30 bp (Fig. 3f), consistent with the reduced sensitivity of short reads for larger insertions reported by Kosugi and Terao [24]. For studies in which the genes of interest lie in long or highly similar duplications, including several medically relevant genes [12], LR sequencing or targeted LR follow-up is therefore likely to be needed, whereas SR sequencing remains adequate for most of the genome. The choice of caller also mattered within each platform. In LowMap, GATK retained higher SR recall at the cost of precision, whereas DeepVariant and Strelka2 with-held more calls (Fig. 2), and on HiFi data, NanoCaller was less accurate than DeepVariant and Clair3 across all stratifications, particularly for INDELs.

Several limitations should be noted. First, not all difficult regions showed a clear difference between platforms. SR SNP calling was largely unaffected in the MHC (Fig. 2). The difficulty of the MHC may arise more from its high polymorphism than from paralog similarity, but our analyses were not designed to test this. Second, the contribution of individual sequence features remains unresolved. Interval length, distance from the interval ends, paralog identity, and copy number were each associated with SR recall, but these features are not independent: larger duplications contained more copies (Fig. 6f,g), and the interior of a long interval may also be where paralog identity is highest. Because each feature was analyzed separately, we cannot determine which of them directly limits SR calling. Third, accuracy could be evaluated only within the GIAB benchmark regions. The most complex parts of segmental duplications and the MHC are partly excluded from these regions, so the differences we observed may underestimate those in the genome as a whole [21].

Our results suggest that SR calling in difficxsxsxsult regions could be improved by adding information that a single short read lacks. Because SR misses were concentrated at positions where reads had a MAPQ of 0, methods that use PSVs to assign such reads to the correct copy, as DuploMap does for long reads [30] and as Parascopy does for paralog-specific copy number estimation from short reads [31], could recover part of the variants currently lost in LowMap and SegDup. Pangenome references, which represent alternative haplotypes and reduce reference bias, are expected to benefit SR alignment most in polymorphic and duplicated regions [21, 29]. In the longer term, complete reference genomes and assembly-based benchmarks will allow evaluation of the regions that current benchmarks leave out [21, 28]. Until such approaches are widely adopted, the length and paralog identity of a region offer a practical way to identify where SR calls are least reliable and where LR data are most needed.

## Methods

### Sequencing data and variant calling

Table 2 summarizes the sequencing datasets and variant-calling tools used in this study. We analyzed whole-genome sequencing (WGS) data from the Genome in a Bottle (GIAB) reference sample HG002 (NA24385) [32], comprising Illumina paired-end short reads and PacBio HiFi long reads. Short reads were aligned to the GRCh38 no-alt analysis set using BWA-MEM (v0.7.18) [33], whereas HiFi reads were aligned using minimap2 (v2.22) [34] with the HiFi preset. Alignments were coordinate-sorted with SAMtools (v1.12). The alignment commands were:

**Table 2.** Variant callers, sequencing datasets and reference resources used in this study.

| Variant Caller | version | Source | Link |
|---|---|---|---|
| NanoCaller | v3.6.0 | Ahsan MU et al. 2021 | https://github.com/WGLab/NanoCaller |
| Clair3 | v1.2.0 | Zheng, Z. et al. 2022 | https://github.com/HKU-BAL/Clair3 |
| DeepVariant | v1.9.0 | Poplin, R. et al. 2018 | https://github.com/google/deepvariant |
| GATK | v4.6.1.0 | McKenna et al. 2010 | https://github.com/broadinstitute/gatk |
| Strelka2 | v2.9.10 | Kim, S. et al. 2018 | https://github.com/Illumina/strelka |
| Sample | Abbreviation | coverage | Source |
| NA24385 | HiFi_L1 | 56x | https://ftp-trace.ncbi.nlm.nih.gov/ReferenceSamples/giab/data/AshkenazimTrio/HG002_NA24385_son/PacBio_CCS_15kb_20kb_chemistry2/reads/ |
| NA24385 | SR_L1 | 74x | https://ftp-trace.ncbi.nlm.nih.gov/ReferenceSamples/giab/data/AshkenazimTrio/HG002_NA24385_son/NIST_Illumina_2x250bps/reads/ |
| NA12878 | NA2878_HiFi | 35x | s3://platinum-pedigree-data/data/hifi/mapped/GRCh38/NA12878.GRCh38.haplotagged.bam |
| NA12878 | NA2878_SR | 30x | s3://platinum-pedigree-data/data/illumina/mapped/GRCh38/2188.v4.2.4.grc38.bam |
| Data |  | Source |  |
| GRCh38 no alt Reference |  | https://storage.googleapis.com/brain-genomics-public/research/platinum/reference/GRCh38.no_alt_analysis_set.fa.gz |  |
| Genome stratification v3.6 |  | https://ftp-trace.ncbi.nlm.nih.gov/ReferenceSamples/giab/release/genome-stratifications/v3.6/ |  |
| NIST v4.2.1 Benchmark |  | https://ftp-trace.ncbi.nlm.nih.gov/ReferenceSamples/giab/release/AshkenazimTrio/HG002_NA24385_son/NISTv4.2.1/ |  |
| Platinum Pedigree Benchmark |  | s3://platinum-pedigree-data/truthset_v1.2/ |  |

~~~
# Short reads (SR_L1)
bwa mem -t 24 -R ‘@RG\tID:D00360\tSM:HG002\tLB:TruSeq\tPL:
   ILLUMINA’ \
  GRCh38.no_alt_analysis_set.fa SR_L1_R1.fastq SR_L1_R2.fastq
\
| samtools sort -@4 -o SR_L1.bam
# HiFi reads (HiFi_L1)
minimap2 -t 30 --MD -Y -L -a -x map-hifi \
  GRCh38.no_alt_analysis_set.fa NA24385_Merged.fastq \
| samtools sort -o HiFi_L1.bam
~~~

Single-nucleotide polymorphisms (SNPs) and small insertions and deletions (INDELs) were called from the short-read alignments using GATK Haplotype-Caller [6], DeepVariant [7], and Strelka2 [35]. For HiFi data, DeepVariant, Clair3 [20], and NanoCaller [36] were used for SNP and INDEL calling. All callers were run on the whole genome with default parameters unless stated otherwise: DeepVariant (v1.9.0) with the `WGS` model for short reads and the `PACBIO` model for HiFi reads; GATK HaplotypeCaller (v4.6.1.0) separately for each of chromosomes 1–22, with the per-chromosome VCFs concatenated with BCFtools (v1.22) [37] and no variant quality score recalibration or hard filtering applied; Strelka2 (v2.9.10) with its germline workflow; Clair3 (v1.2.0) with the `hifi sequel2` model; and NanoCaller (v3.6.0) with the `CCS-HG002` SNP and INDEL models. Variant calls that passed each caller’s own filters were used for evaluation. The commands were ($*{*`REF`*}*, GRCh38 no-alt analysis set; $*{*`BAM`*}*, input alignment):

~~~
# DeepVariant v1.9.0 (MODEL=WGS for SR_L1, MODEL=PACBIO for
   HiFi_L1)
run_deepvariant --model_type=${MODEL} --ref=${REF} --reads=${
   BAM} \
  --output_vcf=output.vcf.gz --output_gvcf=output.g.vcf.gz –
     num_shards=64
# GATK HaplotypeCaller v4.6.1.0, one job per chromosome (i =
   1..22)
gatk --java-options “-Xmx8G” HaplotypeCaller -R ${REF} -I ${BAM
   } \
   -O SR_L1_GATK_chr${i}.vcf -L chr${i}
bcftools concat SR_L1_GATK_chr*.vcf > SR_L1_GATK.vcf
# Strelka2 v2.9.10
configureStrelkaGermlineWorkflow.py --bam ${BAM} --
   referenceFasta ${REF} \
  --runDir workdir
workdir/runWorkflow.py -m local -j 32
# Clair3 v1.2.0
run_clair3.sh --bam_fn=${BAM} --ref_fn=${REF} --threads=20 –
  platform=hifi \
--model_path=/opt/models/hifi_sequel2 --output=${OUT}
# NanoCaller v3.6.0
NanoCaller --bam ${BAM} --ref ${REF} --seq pacbio --snp_model
   CCS-HG002 \
  --indel_model CCS-HG002 --sample NA24385 --cpu 20
~~~

### Evaluation of calling accuracy

Each call set was evaluated in three ways: in the genome-wide benchmark region and five difficult genomic contexts, across sequencing depths, and across INDEL sizes. All three evaluations used the same truth set and regions.

The truth set was the GIAB HG002 GRCh38 v4.2.1 small-variant benchmark for chromosomes 1–22 [25] (Table 2). Genome-wide evaluation was restricted to the benchmark regions released with this truth set, referred to here as AllConf. Five difficult genomic contexts were taken from the GIAB genome stratifications v3.6 [10]: all difficult regions combined (AllD-ifficult), low-mappability regions in which short reads cannot be uniquely aligned (LowMap), the major histocompatibility complex (MHC), segmental duplications (SegDup), and tandem repeats and homopolymers extended by 5 bp on each side (TR/Homo). Only the part of each context lying within AllConf was evaluated, and this overlap was obtained with BEDTools (v2.31.0) [38]:

~~~
bedtools intersect -a HG002_GRCh38_1_22_v4.2.1
   _benchmark_noinconsistent.bed \
  -b ${STRATIFICATION}.bed > ${STRATIFICATION}_highconf.bed
~~~

Variant calls were compared against the truth set using hap.py (v0.3.15) [26] with the `xcmp` comparison engine. Following the hap.py default preprocessing, query variants were decomposed into primitive alleles and INDELs were left-shifted before comparison, while the truth set was used in its original representation. Each call set was evaluated separately within each genomic region, with the BED file of that region supplied as the confident-region file. Recall, precision, and F1 scores for SNPs and INDELs were obtained from the hap.py summary statistics using variants that passed the respective caller’s filtering criteria (Table 1). The command was:

~~~
hap.py HG002_GRCh38_1_22_v4.2.1_benchmark.vcf.gz ${QUERY_VCF} \
  -f ${REGION}.bed -r ${REF} -o happy_results/${REGION}/${
         TOOL} --threads 24
~~~

To assess performance across INDEL sizes, we used the true-positive (TP), false-negative (FN), and false-positive (FP) classifications assigned by hap.py within the AllConf regions. INDEL size was defined as the difference between the alternate and reference allele lengths, with negative values indicating deletions and positive values indicating insertions. For multiallelic sites containing two distinct alternate alleles, the allele with the largest absolute length difference was used. Insertions and deletions were analyzed separately and stratified into 11 size bins: 1, 2, 3, 4, 5–6, 7–8, 9–10, 11– 15, 16–20, 21–30, and ≥ 31 bp. Within each size bin, recall was calculated based on benchmark INDELs, precision based on called INDELs, and the F1 score as their harmonic mean. Size bins containing fewer than 30 benchmark INDELs were excluded from the analysis.

### Subsampling

To assess the effect of sequencing depth, the full-depth SR L1 and HiFi L1 alignments were randomly down-sampled to mean coverages of 10×, 20×, 30×, 40×, and 50× with rasusa (v3.0.0) [39], and the subsampled alignments were processed in the same way as the full-depth alignments (GATK, DeepVariant, and Strelka2 for SR L1; DeepVariant, Clair3, and NanoCaller for HiFi L1), and evaluated in AllConf and AllDifficult. The command was (N = 10, 20, 30, 40, 50):

~~~
rasusa aln --coverage ${N} SR_L1.bam | samtools sort -o SR_L1_$
   {N}x.bam
rasusa aln --coverage ${N} HiFi_L1.bam | samtools sort -o
   HiFi_L1_${N}x.bam
~~~

### Read-level alignment ambiguity

Alignment ambiguity in LowMap was quantified as the proportion of alignments with MAPQ=0, indicating that the aligner could not confidently distinguish between multiple candidate mapping locations. We compared this proportion between short-read and HiFi sequencing data for two groups of variants: variants detected by each platform and variants detected exclusively by one platform.

For the first analysis, we examined DeepVariant calls that passed filtering and fell within LowMap regions for each sequencing platform. At each SNP or INDEL position, overlapping alignments were retrieved from the corresponding BAM file using pysam [40], excluding unmapped and secondary alignments. The proportion of MAPQ 0 alignments was calculated relative to the total number of primary and supplementary alignments overlapping each position. Secondary alignments were excluded to account for differences in alignment reporting between minimap2 and BWA-MEM, particularly the assignment of MAPQ 0 to secondary alignments by minimap2. Variant positions without overlapping mapped alignments were excluded from the analysis.

For the second analysis, we identified platform-specific variants by comparing the DeepVariant call sets within LowMap regions. Variants were considered shared between platforms only when their chromosome, position, reference allele, alternate allele, and variant type matched exactly. For variants detected exclusively in the HiFi data, alignment ambiguity was evaluated using short-read alignments overlapping the corresponding genomic positions. Conversely, HiFi alignments were examined at positions of variants detected exclusively in the short-read data. In both cases, the proportion of MAPQ 0 alignments was calculated using all mapped, non-duplicate alignments at each position, regardless of whether a variant was called by the platform being evaluated.

### Sequence context within difficult regions

Five features of the sequence context were examined. Tract length and distance to the tract edge were related to calling performance and to the proportion of MAPQ 0 reads in both LowMap and SegDup. Sequence identity to the closest paralog, copy number, and duplication size were related to recall and precision in SegDup.

To relate calling performance to tract length and to distance to the tract edge, overlapping or directly adjacent intervals in the LowMap and SegDup stratifications [10] were merged separately into LowMap and SegDup tracts. The merged tracts were not clipped to AllConf, so that tract length and distances reflected the full extent of each tract, whereas only variants within AllConf were evaluated. This analysis used the hap.py [26] output for the full-depth short-read and HiFi DeepVariant callsets. Each benchmark (TRUTH) record labelled TP or FN and each called (QUERY) record labelled TP or FP by hap.py was assigned to the tract containing it. For each record, tract length was the length of that tract, and distance to the tract edge was the distance from the variant position to the nearer end of the tract.

Tract lengths were grouped into seven ranges (≤200, 201–500, 501–1,000, 1,001–2,000, 2,001–5,000, 5,001– 10,000, and *>*10,000 bp), and distances into seven ranges (0–50, 51–125, 126–250, 251–500, 501–1,000, 1,001–2,500, and *>*2,500 bp). For SNPs and INDELs separately, recall was calculated from TRUTH records, precision from QUERY records, and the F1 score from the two, first for each tract-length range and then for each combination of tract-length and distance ranges. Recall and precision were reported only where the corresponding denominator (TP + FN or TP + FP) contained at least 50 records.

For the MAPQ analysis, LowMap and SegDup tracts were constructed in the same way on chromosomes 1–22, X, and Y. Within each tract, sampling points were placed at fixed distances of 0 bp to 100 kb from both ends toward the center. At each point, up to 60 reads were selected at random after unmapped, secondary, supplementary, duplicate, and QC-failed reads were removed, and no read was used more than once. For each read, the distance from its midpoint to the nearer end of the tract was recorded. The proportion of MAPQ 0 reads was then calculated separately for the short-read and HiFi alignments for each combination of the tract-length and distance ranges defined above. Combinations with fewer than 100 reads were not reported.

To estimate paralog similarity and copy number, SegDup intervals on chromosomes 1–22, X, and Y were split into nonoverlapping windows of 150 bp, 1 kb, and 10 kb, and windows with more than 10% N bases were removed. Each window was aligned back to GRCh38 with minimap2 (v2.30) [34], using the short-read preset for 150-bp windows and the assembly-to-reference preset (asm10) for 1-kb and 10-kb windows, and up to 50 secondary alignments were kept per window. The alignment to the window’s own position was discarded, and the remaining alignments that covered at least half of the window were taken as paralogous copies. Identity was calculated as the number of matching bases divided by the alignment block length, and the highest value among the copies was used as the window’s identity to its closest paralog.

Each benchmark variant was assigned to the window containing it. Variants in windows with at least one paralogous copy were grouped by identity (*<*90%, 90–95%, 95–98%, 98–99%, 99–99.5%, 99.5–99.9%, and ≥ 99.9%), and recall in each group was calculated from the hap.py results for the short-read and HiFi Deep-Variant callsets. For 1-kb windows, copy number was defined as the number of other locations to which the window aligned, with alignments on the same chromosome within 1 kb of one another counted as a single location. Copy number was grouped into nine ranges (0, 1, 2, 3, 4–5, 6–9, 10–19, 20–49, and ≥ 50), and recall and precision in each range were reported with 95% Wilson confidence intervals [41]. To relate copy number to duplication size, each 1-kb window was assigned to the SegDup interval containing its midpoint, and the copy number distribution was summarized for each interval size range (1–2, 2–5, 5–10, 10–20, 20–50, 50–100, 100–200, 200–500, and *>*500 kb) by its mean, median, and interquartile range and by the proportion of windows with a copy number of at least 1, 2, 5, and 10.

### Distance to paralogous sequence variants

Paralogous sequence variants (PSVs) are positions at which paralogous copies of a segmental duplication differ in the reference genome. We used the precomputed hg38 PSV database released with DuploMap (v0.9, “All” set) [30]. The database was generated from the reference genome alone. DuploMap was not run on the sequencing data, and the same PSVs were used for both platforms. To build the database, DuploMap takes pairs of homologous copies from the UCSC segmental duplication annotation [42], retaining pairs at least 5 kb long with at least 97% identity, and aligns each pair with minimap2 [34]. It then partitions each alignment at anchors, defined as 6-mers that occur only once within a local neighbourhood of both copies. Each segment between consecutive anchors in which the copies differ is recorded as a PSV, with its coordinates on both copies. Alignment segments with less than 95% similarity do not contribute PSVs. PSVs in low-complexity sequence are retained in the database, and we applied no further filter. PSV records were checked against the GRCh38 no-alt reference, and more than 99% were retained.

Variants in the short-read and HiFi DeepVariant callsets were first matched to the benchmark with hap.py on the complete callsets. Counts for passing calls were then recomputed within AllConf ∩ SegDup, restricted to positions covered by DuploMap alignments that could be assigned to a single duplicated copy (38.9 Mb). For each benchmark (TRUTH) and called (QUERY) variant *v*, the distance to the nearest PSV was defined as

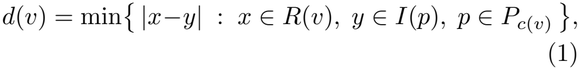

where *c*(*v*) is the duplicated copy containing *v, P*_*c*_ is the set of PSVs on copy *c*, and *R*(*v*) and *I*(*p*) are the reference positions altered by the variant and by the PSV, respectively. An insertion was placed at its left anchor base, a deletion was represented by its deleted bases, and *d*(*v*) = 0 when a variant overlapped a PSV. Variants were normalized with BCFtools norm [37] for this calculation only. The search included PSVs on the same copy that lay outside the evaluation region, but did not extend to other copies. Variants that could not be assigned to a single copy, or whose copy carried no PSV, were excluded (0.08% of benchmark variants). Variants were grouped into seven distance ranges (0, 1–50, 51–150, 151–300, 301–1,000, 1,001–3,000 and *>*3,000 bp). Within each range, recall was calculated from the benchmark-side hap.py labels and precision from the call-side labels. Read and fragment lengths were estimated from alignments sampled in segmental duplications. The short reads were 2 × 250 bp, with a median fragment length of 404 bp, and the HiFi reads had a median length of 13.5 kb.

## Data availability

PacBio HiFi and Illumina NA24385 sequencing reads are available at NCBI. Benchmarking call set and high-confidence regions for NA24385 were obtained from NIST v4.2.1 benchmark dataset. NA12878 sequencing reads, benchmarking call set and high-confidence regions are available under Platinume Pedigree project’s S3 repository [43]. The reference genome used in the study is GRCh38 no alt andlysis set. Specific links for variance callers and aforementioned data are listed in Table 2

## Code availability

All code and related files are available at https://github.com/maiziezhoulab/Benchmark_small_variants_shortreads_vs_longreads.

## References

1. 1000 Genomes Project Consortium: A global reference for human genetic variation. Nature 526, 68–74 (2015). doi:10.1038/nature15393

2. Mullaney, J.M., Mills, R.E., Pittard, W.S., Devine, S.E.: Small insertions and deletions (INDELs) in human genomes. Human Molecular Genetics 19(R2), 131–136 (2010). doi:10.1093/hmg/ddq400

3. Montgomery, S.B., Goode, D.L., Kvikstad, E., et al.: The origin, evolution, and functional impact of short insertion–deletion variants identified in 179 human genomes. Genome Research 23(5), 749–761 (2013). doi:10.1101/gr.148718.112

4. van Dijk, E.L., Auger, H., Jaszczyszyn, Y., Thermes, C.: Ten years of next-generation sequencing technology. Trends in Genetics 30(9), 418–426 (2014). doi:10.1016/j.tig.2014.07.001

5. Bentley, D.R., Balasubramanian, S., Swerdlow, H.P., et al.: Accurate whole human genome sequencing using reversible terminator chemistry. Nature 456, 53–59 (2008). doi:10.1038/nature07517

6. DePristo, M.A., Banks, E., Poplin, R., et al.: A framework for variation discovery and genotyping using next-generation DNA sequencing data. Nature Genetics 43(5), 491–498 (2011). doi:10.1038/ng.806

7. Poplin, R., Chang, P.-C., Alexander, D., et al.: A universal SNP and small-indel variant caller using deep neural networks. Nature Biotechnology 36(10), 983–987 (2018). doi:10.1038/nbt.4235

8. Olson, N.D., Wagner, J., McDaniel, J., et al.: PrecisionFDA Truth Challenge V2: calling variants from short and long reads in difficult-to-map regions. Cell Genomics 2(5), 100129 (2022). doi:10.1016/j.xgen.2022.100129

9. Ebbert, M.T.W., Jensen, T.D., Jansen-West, K., et al.: Systematic analysis of dark and camouflaged genes reveals disease-relevant genes hiding in plain sight. Genome Biology 20, 97 (2019). doi:10.1186/s13059-019-1707-2

10. Dwarshuis, N., Kalra, D., McDaniel, J., et al.: The GIAB genomic stratifications resource for human reference genomes. Nature Communications 15, 9029 (2024). doi:10.1038/s41467-024-53260-y

11. Mandelker, D., Schmidt, R.J., Ankala, A., et al.: Navigating highly homologous genes in a molecular diagnostic setting: a resource for clinical next-generation sequencing. Genetics in Medicine 18, 1282–1289 (2016). doi:10.1038/gim.2016.58

12. Wagner, J., Olson, N.D., Harris, L., et al.: Curated variation benchmarks for challenging medically relevant autosomal genes. Nature Biotechnology 40, 672–680 (2022). doi:10.1038/s41587-021-01158-1

13. Mantere, T., Kersten, S., Hoischen, A.: Long-read sequencing emerging in medical genetics. Frontiers in Genetics 10, 426 (2019). doi:10.3389/fgene.2019.00426

14. Ameur, A., Kloosterman, W.P., Hestand, M.S.: Single-molecule sequencing: towards clinical applications. Trends in Biotechnology 37(1), 72–85 (2019). doi:10.1016/j.tibtech.2018.07.013

15. Eid, J., Fehr, A., Gray, J., et al.: Real-time DNA sequencing from single polymerase molecules. Science 323(5910), 133–138 (2009). doi:10.1126/science.1162986

16. Wenger, A.M., Peluso, P., Rowell, W.J., et al.: Accurate circular consensus long-read sequencing improves variant detection and assembly of a human genome. Nature Biotechnology 37, 1155–1162 (2019). doi:10.1038/s41587-019-0217-9

17. Branton, D., Deamer, D.W., Marziali, A., et al.: The potential and challenges of nanopore sequencing. Nature Biotechnology 26(10), 1146–1153 (2008). doi:10.1038/nbt.1495

18. Jain, M., Koren, S., Miga, K.H., et al.: Nanopore sequencing and assembly of a human genome with ultra-long reads. Nature Biotechnology 36, 338–345 (2018). doi:10.1038/nbt.4060

19. Shafin, K., Pesout, T., Chang, P.-C., et al.: Haplotype-aware variant calling with PEPPER-Margin-DeepVariant enables high accuracy in nanopore long-reads. Nature Methods 18, 1322–1332 (2021). doi:10.1038/s41592-021-01299-w

20. Zheng, Z., Li, S., Su, J., Leung, A.W.-S., Lam, T.-W., Luo, R.: Symphonizing pileup and full-alignment for deep learning-based long-read variant calling. Nature Computational Science 2(12), 797–803 (2022). doi:10.1038/s43588-022-00387-x

21. Zheng, Z., Sedlazeck, F.J., Luo, R.: Small variant calling in the era of long-read sequencing. Nature Reviews Genetics (2026). doi:10.1038/s41576-026-01020-7

22. Zook, J.M., Hansen, N.F., Olson, N.D., et al.: A robust benchmark for detection of germline large deletions and insertions. Nature Biotechnology 38, 1347–1355 (2020). doi:10.1038/s41587-020-0538-8

23. Ebert, P., Audano, P.A., Zhu, Q., et al.: Haplotype-resolved diverse human genomes and integrated analysis of structural variation. Science 372(6537), 7117 (2021). doi:10.1126/science.abf7117

24. Kosugi, S., Terao, C.: Comparative evaluation of SNVs, indels, and structural variations detected with short- and long-read sequencing data. Human Genome Variation 11, 1–10 (2024). doi:10.1038/s41439-024-00276-x

25. Wagner, J., Olson, N.D., Harris, L., et al.: Benchmarking challenging small variants with linked and long reads. Cell Genomics 2(5), 100128 (2022). doi:10.1016/j.xgen.2022.100128

26. Krusche, P., Trigg, L., Boutros, P.C., et al.: Best practices for benchmarking germline small-variant calls in human genomes. Nature Biotechnology 37(5), 555–560 (2019). doi:10.1038/s41587-019-0054-x

27. Cleary, J.G., Braithwaite, R., Gaastra, K., et al.: Comparing variant call files for performance benchmarking of next-generation sequencing variant calling pipelines. bioRxiv 023754 (2015). doi:10.1101/023754

28. Nurk, S., Koren, S., Rhie, A., et al.: The complete sequence of a human genome. Science 376(6588), 44–53 (2022). doi:10.1126/science.abj6987

29. Liao, W.-W., Asri, M., Ebler, J., et al.: A draft human pangenome reference. Nature 617, 312–324 (2023). doi:10.1038/s41586-023-05896-x

30. Prodanov, T., Bansal, V.: Sensitive alignment using paralogous sequence variants improves long-read mapping and variant calling in segmental duplications. Nucleic Acids Research 48(19), 114 (2020). doi:10.1093/nar/gkaa829

31. Prodanov, T., Bansal, V.: Robust and accurate estimation of paralog-specific copy number for duplicated genes using whole-genome sequencing. Nature Communications 13, 3221 (2022). doi:10.1038/s41467-022-30930-3

32. Zook, J.M., Catoe, D., McDaniel, J., et al.: Extensive sequencing of seven human genomes to characterize benchmark reference materials. Scientific Data 3, 160025 (2016). doi:10.1038/sdata.2016.25

33. Li, H.: Aligning sequence reads, clone sequences and assembly contigs with BWA-MEM. arXiv:1303.3997 (2013). 1303.3997

34. Li, H.: Minimap2: pairwise alignment for nucleotide sequences. Bioinformatics 34(18), 3094–3100 (2018). doi:10.1093/bioinformatics/bty191

35. Kim, S., Scheffler, K., Halpern, A.L., et al.: Strelka2: fast and accurate calling of germline and somatic variants. Nature Methods 15(8), 591–594 (2018). doi:10.1038/s41592-018-0051-x

36. Ahsan, M.U., Liu, Q., Fang, L., Wang, K.: NanoCaller for accurate detection of SNPs and indels in difficult-to-map regions from long-read sequencing by haplotype-aware deep neural networks. Genome Biology 22, 261 (2021). doi:10.1186/s13059-021-02472-2

37. Danecek, P., Bonfield, J.K., Liddle, J., Marshall, J., Ohan, V., Pollard, M.O., Whitwham, A., Keane, T., McCarthy, S.A., Davies, R.M., Li, H.: Twelve years of SAMtools and BCFtools. GigaScience 10(2), 008 (2021). doi:10.1093/gigascience/giab008

38. Quinlan, A.R., Hall, I.M.: BEDTools: a flexible suite of utilities for comparing genomic features. Bioinformatics 26(6), 841–842 (2010). doi:10.1093/bioinformatics/btq033

39. Hall, M.B.: Rasusa: Randomly subsample sequencing reads to a specified coverage. Journal of Open Source Software 7(69), 3941 (2022). doi:10.21105/joss.03941

40. Pysam developers: pysam: a Python module for reading and manipulating SAM/BAM/VCF/BCF files. https://github.com/pysam-developers/pysam. Accessed 14 November 2026

41. Wilson, E.B.: Probable inference, the law of succession, and statistical inference. Journal of the American Statistical Association 22(158), 209–212 (1927). doi:10.1080/01621459.1927.10502953

42. Bailey, J.A., Gu, Z., Clark, R.A., Reinert, K., Samonte, R.V., Schwartz, S., Adams, M.D., Myers, E.W., Li, P.W., Eichler, E.E.: Recent segmental duplications in the human genome. Science 297(5583), 1003–1007 (2002). doi:10.1126/science.1072047

43. Kronenberg, Z., Nolan, C., Porubsky, D., Mokveld, T., Rowell, W.J., Lee, S., Dolzhenko, E., Chang, P.-C., Holt, J.M., Saunders, C.T., et al.: The platinum pedigree: a long-read benchmark for genetic variants. Nature Methods 22(8), 1669–1676 (2025)

