## Supplementary Information for "A stratified benchmark reveals distinct short- and long-read performance gradients within difficult genomic regions"

Zhang et al.

#### Contents

**1 Supplementary Tables** **2**
1 Small-variant calling on NA12878 . . . . . . . . . . . . . . . . . . . . 2

**2 Supplementary Figures** **3**
1 NA12878 radar plot . . . . . . . . . . . . . . . . . . . . . . . . . . . 4
2 Calling performance decrease in large LowMap and SegDup regions 5

### 1 Supplementary Tables

| INDEL F1 | Clair3 | DeepVariant | NanoCaller |
|---|---|---|---|
| AllConf | 94.4% | 98.4% | 90.2% |
| AllDiff | 92.8% | 98.0% | 89.5% |
| LowMap | 89.2% | 93.9% | 82.9% |
| MHC | 95.7% | 98.1% | 91.6% |
| SegDup | 90.5% | 95.3% | 82.9% |
| TR/Homo | 92.5% | 97.9% | 89.5% |
| INDEL Recall | Clair3 | DeepVariant | NanoCaller |
| AllConf | 95.5% | 98.4% | 86.9% |
| AllDiff | 94.3% | 98.0% | 85.9% |
| LowMap | 89.8% | 92.9% | 78.4% |
| MHC | 96.4% | 98.2% | 89.7% |
| SegDup | 91.0% | 94.4% | 79.7% |
| TR/Homo | 94.1% | 98.0% | 85.8% |
| INDEL Precision | Clair3 | DeepVariant | NanoCaller |
| AllConf | 93.2% | 98.3% | 93.8% |
| AllDiff | 91.4% | 97.9% | 93.4% |
| LowMap | 88.6% | 94.9% | 87.9% |
| MHC | 95.0% | 98.0% | 93.7% |
| SegDup | 90.0% | 96.2% | 86.3% |
| TR/Homo | 91.1% | 97.9% | 93.5% |
| SNP F1 | Clair3 | DeepVariant | NanoCaller |
| AllConf | 99.4% | 99.6% | 98.2% |
| AllDiff | 97.7% | 98.2% | 93.6% |
| LowMap | 96.7% | 97.4% | 94.0% |
| MHC | 99.5% | 99.6% | 98.8% |
| SegDup | 94.4% | 96.3% | 89.0% |
| TR/Homo | 96.8% | 97.1% | 89.3% |
| SNP Recall | Clair3 | DeepVariant | NanoCaller |
| AllConf | 99.6% | 99.5% | 99.2% |
| AllDiff | 98.2% | 98.1% | 97.0% |
| LowMap | 96.5% | 97.0% | 96.9% |
| MHC | 99.5% | 99.5% | 99.0% |
| SegDup | 94.8% | 95.5% | 95.3% |
| TR/Homo | 97.8% | 97.2% | 94.6% |
| SNP Precision | Clair3 | DeepVariant | NanoCaller |
| AllConf | 99.3% | 99.6% | 97.2% |
| AllDiff | 97.2% | 98.2% | 90.4% |
| LowMap | 96.9% | 97.9% | 91.4% |
| MHC | 99.6% | 99.6% | 98.5% |
| SegDup | 94.0% | 97.1% | 83.6% |
| TR/Homo | 95.9% | 97.0% | 84.5% |

| INDEL F1 | GATK | DeepVariant | Strelka2 |
|---|---|---|---|
| AllConf | 91.7% | 93.7% | 94.9% |
| AllDiff | 89.8% | 92.0% | 93.5% |
| LowMap | 78.8% | 82.9% | 82.2% |
| MHC | 94.1% | 93.9% | 94.2% |
| SegDup | 83.7% | 88.0% | 88.1% |
| TR/Homo | 89.6% | 91.7% | 93.3% |
| INDEL Recall | GATK | DeepVariant | Strelka2 |
| AllConf | 89.7% | 93.8% | 92.2% |
| AllDiff | 87.6% | 92.1% | 90.2% |
| LowMap | 70.8% | 76.9% | 73.2% |
| MHC | 93.7% | 93.3% | 92.1% |
| SegDup | 78.2% | 84.8% | 82.2% |
| TR/Homo | 87.4% | 92.0% | 90.0% |
| INDEL Precision | GATK | DeepVariant | Strelka2 |
| AllConf | 93.8% | 93.6% | 97.6% |
| AllDiff | 92.2% | 91.9% | 97.0% |
| LowMap | 88.8% | 89.9% | 93.6% |
| MHC | 94.5% | 94.4% | 96.4% |
| SegDup | 90.1% | 91.5% | 95.0% |
| TR/Homo | 91.9% | 91.5% | 96.9% |
| SNP F1 | GATK | DeepVariant | Strelka2 |
| AllConf | 97.3% | 99.2% | 98.8% |
| AllDiff | 93.4% | 96.4% | 95.1% |
| LowMap | 88.1% | 93.0% | 89.8% |
| MHC | 99.2% | 99.3% | 99.2% |
| SegDup | 87.0% | 92.1% | 89.6% |
| TR/Homo | 92.7% | 95.4% | 94.1% |
| SNP Recall | GATK | DeepVariant | Strelka2 |
| AllConf | 95.6% | 98.8% | 98.1% |
| AllDiff | 90.4% | 95.1% | 92.1% |
| LowMap | 81.7% | 89.8% | 82.6% |
| MHC | 99.2% | 99.0% | 98.8% |
| SegDup | 81.2% | 88.2% | 82.2% |
| TR/Homo | 90.9% | 94.4% | 92.0% |
| SNP Precision | GATK | DeepVariant | Strelka2 |
| AllConf | 99.1% | 99.5% | 99.5% |
| AllDiff | 96.5% | 97.8% | 98.2% |
| LowMap | 95.6% | 96.5% | 98.5% |
| MHC | 99.3% | 99.5% | 99.6% |
| SegDup | 93.6% | 96.3% | 98.4% |
| TR/Homo | 94.5% | 96.3% | 96.4% |

**Supplementary Table 1:** Precision, recall and F1 score of HiFi (NA12878_HiFi) and short-read (NA12878_SR) variant callers across genomic stratifications in NA12878, evaluated with hap.py against the Platinum Pedigree truth set v1.2 benchmark.

#### 2 Supplementary Figures

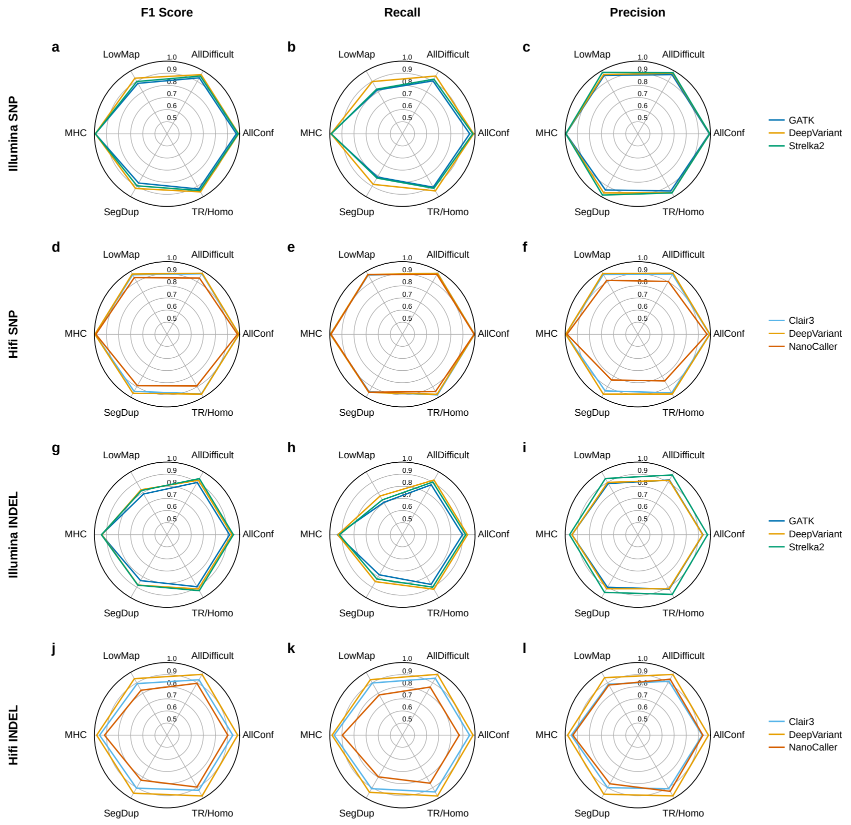


**Supplementary Figure 1: Small-variant calling performance across genomic stratifications.** Radar plots of F1 score (left column), recall (middle column) and precision (right column) for the short-read (NA12878_SR; Illumina) and long-read (NA12878_Hifi; PacBio HiFi) datasets of NA12878, evaluated with hap.py against the Platinum Pedigree truth set v1.2 in six stratifications: AllConf, AllDifficult, LowMap, MHC, SegDup and TR/Homo. Concentric circles mark values from 0.5 to 1.0. **a–c** SR SNPs called with GATK, DeepVariant and Strelka2. **d–f** HiFi SNPs called with Clair3, DeepVariant and NanoCaller. **g–i** SR INDELs called with GATK, DeepVariant and Strelka2. **j–l** HiFi INDELs called with Clair3, DeepVariant and NanoCaller.

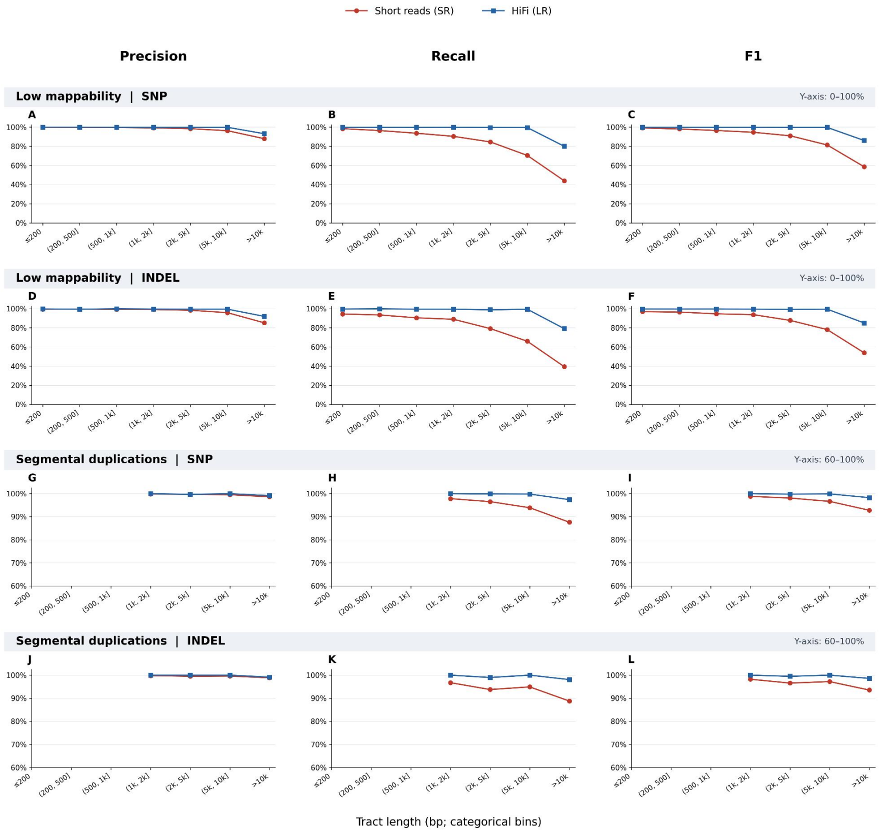


**Supplementary Figure 2: Variant calling performance declines with increasing tract length in low-mappability regions and segmental duplications.** Precision, recall, and F1 scores are compared between Illumina short reads (SR) and PacBio HiFi long reads (LR) across different tract length bins. Both sequencing platforms exhibit reduced calling performance in longer tracts, with a more pronounced decline in recall for SR. **a–c** SNP calling performance across tract length bins in low-mappability regions. **d–f** INDEL calling performance across tract length bins in low-mappability regions. **g–i** SNP calling performance across tract length bins in segmental duplications. **j–l** INDEL calling performance across tract length bins in segmental duplications.
